# Primate lineage specification requires suppression of Alu hyperediting

**DOI:** 10.64898/2026.01.14.699349

**Authors:** Emily J. Park, Yingzhi Cui, Florencia Levin-Ferreyra, Víctor López Soriano, Hao Wu, Núria Lupión-Garcia, Caroline M. Sands, Patrizia Pessina, M. Cecilia Guerra, Jorge Botas, Li-Yu Chen, Katerina Cermakova, H. Courtney Hodges, Lluis Morey, Joshua J. Coon, Jun Wu, Aryeh Warmflash, Eric Van Nostrand, Michael S. Hoetker, Bruno Di Stefano

## Abstract

Understanding human specific mechanisms of cell fate control is essential for advancing developmental biology and regenerative medicine. Here, we identify the ILF2/3 complex as a critical regulator of primate cell fate transitions. Using genetically and epigenetically engineered gastruloids and adult stem cells, we show that ILF2/3 is required for gastrulation in primates but not mice, and for differentiation of adult progenitor cells. Mechanistically, ILF2/3 directly binds Alu elements in chromatin-associated RNAs and shields them from ADAR1-mediated adenosine-to-inosine (A-to-I) editing. Acute ILF2/3 degradation increases A-to-I editing at Alu elements, but not murine retrotransposons, leading to aberrant splicing and nonsense-mediated decay of transcripts encoding key chromatin regulators in primate cells. This in turn destabilizes the epigenetic landscape and blocks lineage commitment across all three germ layers. Re-expression of correctly spliced chromatin regulators rescues differentiation defects in ILF2/3-deficient cells, functionally linking Alu editing control to chromatin regulation and cell fate. These findings define an evolutionary mechanism that restrains retrotransposon-associated RNA editing to preserve proteome integrity and enable primate-specific developmental programs.

## MAIN

How species-specific regulatory mechanisms have emerged to instruct cell fate remains a fundamental question in biology. Cell fate specification depends on the interplay of transcriptional and post-transcriptional regulatory mechanisms. Transcriptional regulation is mediated by transcription factors and chromatin remodelers that orchestrate gene expression and genomic architecture to regulate cell identity^2–4^. Once RNAs are transcribed, post-transcriptional mechanisms regulate cell fate by modulating RNA localization, stability, translation efficiency, and other functions^5–16^. Although core principles of cell fate specification are conserved across mammals^17–20^, genome composition has diverged substantially between species^21–23^, suggesting that lineage-specific regulatory mechanisms have evolved to accommodate these differences.

Crucially, co-transcriptional processes acting on pre-mRNAs, including splicing, polyadenylation, and RNA editing, have emerged as key regulators of this lineage-specific divergence^6, 7, 9, 13, 24–28^. These mechanisms can contribute to the regulation of species-specific genomic elements, such as transposons, and several have been implicated in evolutionary innovation. For example, RNA editing occurs across metazoans but primarily targets evolutionarily younger, more species-restricted repetitive elements in non-coding regions of transcripts^28^. Similarly, in humans, transposable elements harboring alternative splice sites tend to be less conserved and are repressed by splicing regulatory proteins^29,30^. However, how co-transcriptional mechanisms are differentially co-opted across species to control cell fate during developmental transitions remains unclear.

An emerging class of proteins that bind both DNA and RNA (DRBPs) is positioned to facilitate co-transcriptional control and orchestrate species-specific mechanisms by maintaining proximity to chromatin-associated RNAs^8, 31–34^. For example, TDP-43 and KRAB zinc finger proteins bind both DNA and RNA, functioning in transcriptional and post-transcriptional processes like transposable element repression and pre-mRNA splicing^35–39^. However, the broader mechanistic role of DRBPs in co-transcriptional regulation during embryonic cell fate specification remains to be elucidated.

Here, we identify two DRBPs, ILF2 and ILF3, as essential regulators of primate cell fate across developmental contexts and tissue types. The ILF2/3 complex is required for primate, but not mouse, gastrulation and for the differentiation of adult human progenitors across multiple lineages. Mechanistically, ILF2/3 binds and inhibits ADAR1 to suppress RNA editing at primate-specific Alu elements. Acute degradation of ILF2/3 induces rapid RNA hyperediting, leading to aberrant splicing of transcripts encoding chromatin regulators, their nonsense-mediated decay, and disrupted cell fate specification. Our findings establish a mechanistic link between RNA editing and species-specific cell fate, revealing how primates co-opted ILF2/3 as an evolutionary safeguard to constrain Alu-associated transcript instability.

## RESULTS

### Dual DNA- and RNA-binding proteins control primate cell fate

DRBPs play a crucial role in co-transcriptional processes^32, 40^, yet their function in coordinating cell fate remains unexplored. To identify DRBPs controlling human development, we analyzed genome-wide loss-of-function screens for genes required during human pluripotent stem cell (hPSC) differentiation^1^. Given the species-specificity of DRBP functions^32^ and the need to identify human-specific regulators of stem cell plasticity, we excluded genes previously implicated in mouse pluripotency^41–43^. This analysis identified ILF2 and ILF3 as the most enriched candidates uniquely required for human pluripotency exit (**Fig. 1a**). ILF3 heterodimerizes with ILF2^44^ and functions as a transcription factor^45–49^, but recent work has implicated it in RNA metabolism^49–52^. While ILF2 and ILF3 are highly conserved (**Extended Data Fig. 1a**), their transcripts show higher expression and translation in human versus mouse embryos^53–55^ (**Extended Data Fig. 1b,c**). Furthermore, neither protein appeared to be critical for self-renewal or differentiation in mice based on screens performed in mouse embryonic stem cells^43^ (**Extended Data Fig. 1d**). These findings suggest that ILF2 and ILF3 may have evolved functions in regulating human cell fate.

**Figure 1.**
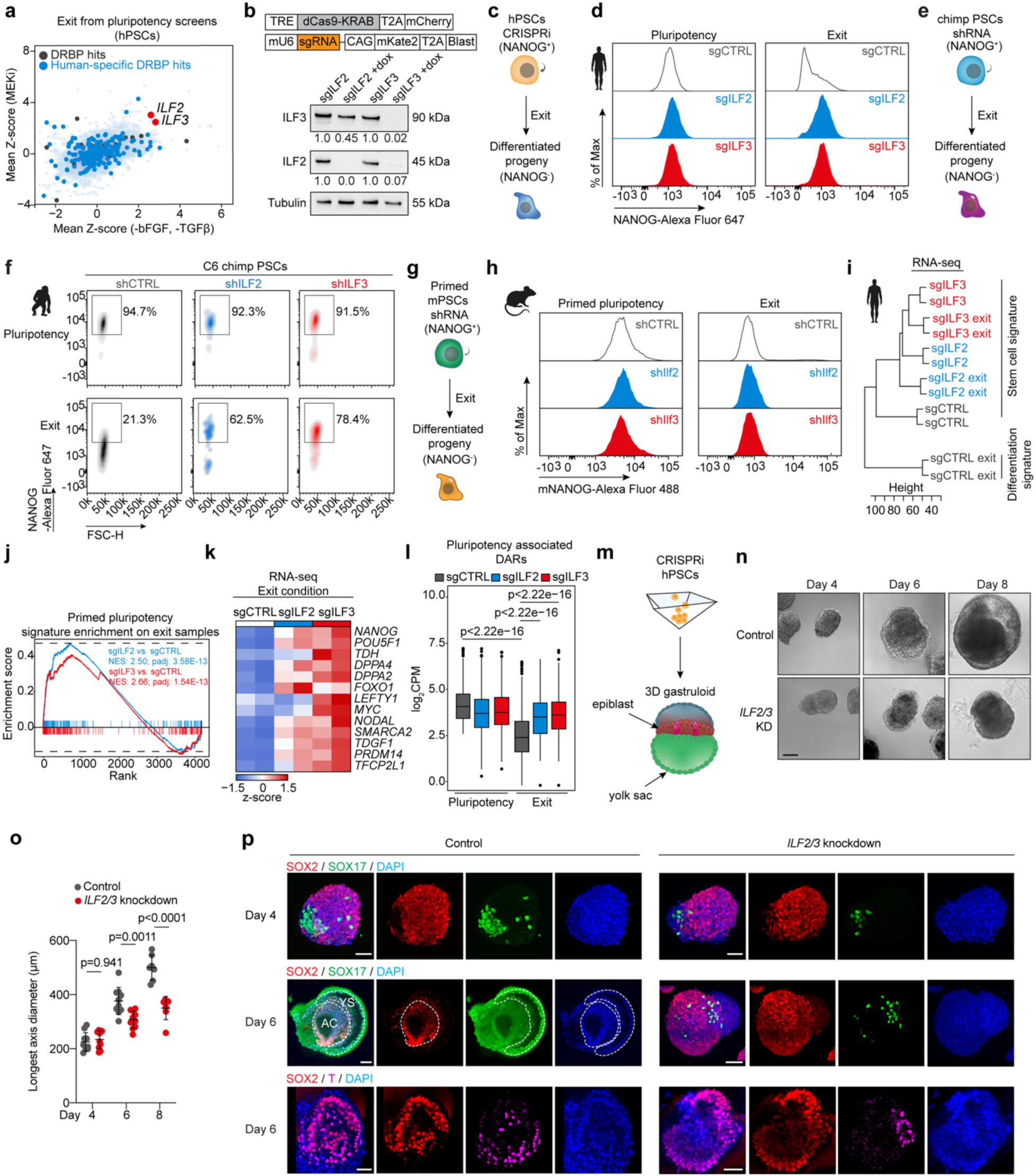
ILF2 and ILF3 are required for human gastrulation. (**a**) Results from genome-wide loss-of-function screens in human pluripotent stem cells (hPSCs) during pluripotency exit induced by either TGFβ and bFGF withdrawal or MAPK pathway inhibition^1^, depicting mean Z-score from three replicates. (**b**) CRISPRi (top) and representative western blot with signal quantification (bottom) depicting validation of *ILF2* and *ILF3* knockdown efficiency after 3 days of doxycycline treatment in hPSCs expressing targeted guide RNAs. (**c**) Experimental design for pluripotency exit assays. (**d**) Representative flow cytometry quantification of NANOG in hPSCs under self-renewal and exit conditions by MAPK pathway inhibition following *ILF2* or *ILF3* knockdown. (**e**) Schematic of chimpanzee (*Pan troglodytes*) PSC differentiation experiments. (**f**) Representative flow cytometry quantification of NANOG-positive chimpanzee PSCs under self-renewal or exit conditions following *ILF2* or *ILF3* knockdown. (**g**) Schematic of mouse ESC differentiation experiments. (**h**) Representative flow cytometry quantification of NANOG-positive mouse ESCs under primed pluripotency or exit conditions following *Ilf2* or *Ilf3* knockdown. (**i**) Hierarchical clustering of RNA-seq datasets from hPSCs in self-renewal and exit conditions. (**j**) Gene Set Enrichment Analysis (GSEA) of pluripotency-associated genes in *ILF2*- or *ILF3*-depleted cells versus control during exit conditions (ILF2: NES=2.50, p=3.58E-13; ILF3: NES=2.66, p= 1.54E-13). (**k**) Differential gene expression analysis comparing control and *ILF2* or *ILF3*-knockdown hPSCs after exit from pluripotency (n = 2 biological replicates; fold change > 1.5; P < 0.05). Red and blue indicate up- and down-regulated genes, respectively. (**l**) Quantification of pluripotency-associated differentially accessible regions in control and *ILF2* or *ILF3*-depleted cells under self-renewal and exit conditions (n = 2 biological replicates, P values calculated using paired Wilcoxon rank-sum test). (**m**) Experimental design for three-dimensional human peri-gastruloid formation. (**n**) Representative brightfield images of *ILF2*- and *ILF3*-depleted human peri-gastruloids. Scale bar, 100 µm. (**o**) Analysis of longest axis length *ILF2*- and *ILF3*-depleted human peri-gastruloids (n = 7-10 biological replicates; mean ± s.d; P values determined by two-way ANOVA with Šidák’s multiple comparisons test). (**p**) Immunofluorescence analysis of human peri-gastruloids showing SOX2 (red), SOX17 (green), T (pink), and nuclear DAPI staining (blue) in control and *ILF2* or *ILF3*-depleted cells. Scale bar, 50 µm.

To investigate the roles of ILF2 and ILF3 during pluripotency exit, we established an inducible CRISPR interference (CRISPRi) system in hPSCs^56, 57^. We first confirmed the nuclear localization of ILF2 and ILF3 through colocalization with DAPI in hPSCs (**Extended Data Fig. 1e**). CRISPRi-mediated silencing effectively depleted both ILF2 and ILF3 at the RNA and protein levels (**Fig. 1b and Extended Data Fig. 1f,g**). Consistent with previous studies^44^, *ILF3* depletion did not suppress *ILF2* transcript levels but caused a marked reduction in ILF2 protein (**Fig. 1b and Extended Data Fig. 1f,g**), indicating that ILF2 stability depends upon the presence of ILF3.

To assess the role of ILF2 and ILF3 in pluripotency exit, we tracked NANOG protein levels during differentiation (**Fig. 1c**). While suppression of *ILF2* or *ILF3* did not affect NANOG expression under self-renewal conditions, it delayed downregulation of NANOG during differentiation induced by TGFβ and bFGF withdrawal or MAPK pathway inhibition ^1, 56^ (**Fig. 1d and Extended Data Fig. 2a,b**). We corroborated these findings using shRNA-mediated knockdown in OCT4-GFP reporter human ESCs^58^ (**Extended Data Fig. 2c**), where depletion of either factor maintained OCT4-GFP and NANOG expression under differentiation conditions (**Extended Data Fig. 2d,e**). These data demonstrate that ILF2 and ILF3 are crucial for exit from pluripotency in hPSCs.

To determine whether the roles of ILF2 and ILF3 are conserved across species, we compared their functions in chimpanzee and mouse. Consistent with our findings in hPSCs, suppression of *ILF2* or *ILF3* in two different chimpanzee PSC lines (C6, C7)^59^ delayed the downregulation of NANOG protein and pluripotency-associated transcripts (**Fig. 1e,f and Extended Data Fig. 2f,g**). To enable cross-species comparison of equivalent pluripotent states, we converted naïve mouse ESCs to epiblast-like cells, which model the primed state^60, 61^. In contrast to our findings in primate PSCs, suppression of *Ilf2* or *Ilf3* in mouse primed ESCs did not impair self-renewal or differentiation (**Fig. 1g,h and Extended Data Fig. 2h,i**). These findings align with previous studies demonstrating that ILF2 and ILF3 are dispensable for gastrulation in zebrafish and mice^62, 63^, highlighting species-specific mechanisms underlying ILF2/3 function in cell fate determination.

Collectively, these findings indicate that ILF2 and ILF3 regulate primate embryonic cell fate through mechanisms not conserved in rodents.

### ILF2 and ILF3 are required for dissolution of the pluripotency transcriptional network

To investigate how ILF2 and ILF3 regulate exit from pluripotency, we analyzed gene expression changes following *ILF2* or *ILF3* silencing under self-renewal and differentiation conditions. Silencing of either factor induced similar transcriptional changes (r^Pluripotency^= 0.87, r^Differentiation^= 0.89) (**Extended Data Fig. 3a,b**). Hierarchical clustering further revealed that *ILF2*- and *ILF3*-depleted cells under differentiation conditions clustered more closely with pluripotent control cells than with their differentiated counterparts (**Fig. 1i**). Accordingly, *ILF2*- and *ILF3*-depleted cells maintained high expression of pluripotency-associated transcripts (e.g., *NANOG*, *POU5F1*) (**Fig. 1j,k**). These findings demonstrate that both ILF2 and ILF3 are essential for the dissolution of the pluripotency transcriptional program.

Since transcriptional rewiring is tightly linked to chromatin remodeling, we assessed whether *ILF2* and *ILF3* influence chromatin accessibility during pluripotency exit. To this end, we performed Assay for Transposase-Accessible Chromatin sequencing (ATAC-seq)^64^ to assess changes in chromatin accessibility in *ILF2*- or *ILF3*-depleted hPSCs under self-renewal and differentiation conditions. We observed thousands of changes in global chromatin accessibility compared to control cells, with strong correlation between *ILF2*- and *ILF3*-depleted conditions (r^Pluripotency^= 0.51, r^Differentiation^= 0.50) (**Extended Data Fig. 3c,d**). Clustering analysis showed that *ILF2*- and *ILF3*-depleted cells under differentiation conditions grouped more closely with undifferentiated cells, with regions that normally become less accessible during differentiation retaining an open chromatin state in *ILF2*- or *ILF3*-depleted cells (**Fig. 1l and Extended Data Fig. 3e**).

Together, our data provide evidence that ILF2 and ILF3 are critical for the dissolution of the pluripotency transcriptional network and the proper remodeling of the epigenetic landscape of differentiated cells.

### ILF2/3 is required for human, but not mouse, gastruloid formation

Given their critical role in dissolving human pluripotency (**Fig. 1c,d and Extended Data Fig. 2a-e**), we hypothesized that ILF2 and ILF3 also regulate germ layer specification. To test this, we employed three-dimensional peri-gastruloid models^65^, which recapitulate key aspects of human peri-gastrulation development (**Fig. 1m**). ILF2/3-depleted peri-gastruloids were significantly smaller than controls throughout differentiation, consistent with a differentiation defect (**Fig. 1n,o**). By day 6, ILF2/3-depleted peri-gastruloids exhibited significant reduction in SOX17+ cells and mesendoderm markers, while the epiblast marker SOX2 was retained, indicating dysregulated germ layer specification (**Fig. 1p and Extended Data Fig. 3f**). We corroborated these findings using a 2D gastrulation model^66^, which showed that ILF2/3 loss disrupted lineage specification (**Extended Data Fig. 3g,h**). These data establish the ILF2/3 complex as a key regulator of human gastrulation, where depletion perturbs germ layer specification across multiple experimental systems.

We next examined whether ILF2 and ILF3 play a similar role in mouse gastrulation^67, 68^ (**Extended Data Fig. 3i**). Consistent with our observations in mouse ESC differentiation assays (**Fig. 1g,h**), suppression of *Ilf2* or *Ilf3* in mouse gastruloids had no significant effect on morphology or lineage marker expression compared to controls (**Extended Data Fig. 3i-l**). While standard mouse gastruloids reflect different developmental dynamics than their human counterparts, the consistent absence of phenotypes across multiple murine platforms—including ESC differentiation assays, gastruloids, and *Ilf3* knockout mice^63^—contrasts with the robust effects observed across human pluripotency and gastrulation models, supporting a species-specific role for ILF2 and ILF3 in primate embryonic development.

### ILF2 and ILF3 direct adult stem and progenitor cell differentiation

To investigate whether ILF2 and ILF3 act as general safeguards of human stem cell identity across lineages, we tested the effects of *ILF2* or *ILF3* knockdown in neural progenitor cells (NPCs) undergoing differentiation into neurons. While control NPCs differentiated efficiently into TUJ1-expressing neurons, suppression of *ILF2* or *ILF3* significantly impaired neuronal differentiation by day 6 (**Fig. 2a,b and Extended Data Fig. 4a**). *ILF2*- and *ILF3*-depleted NPCs also exhibited upregulation of proliferation genes and downregulation of genes associated with neuronal differentiation (**Fig. 2c and Extended Data Fig. 4b,c**). For example, following *ILF2* and *ILF3* suppression, the neuronal master regulators *ASCL1*^69–71^ and *NEUROD4*^72, 73^ were significantly downregulated (**Fig. 2d**), consistent with a failure to properly activate a neuron-specific gene signature. These findings parallel our observations in PSCs and highlight the important roles of ILF2 and ILF3 in facilitating NPC-to-neuron differentiation.

**Figure 2.**
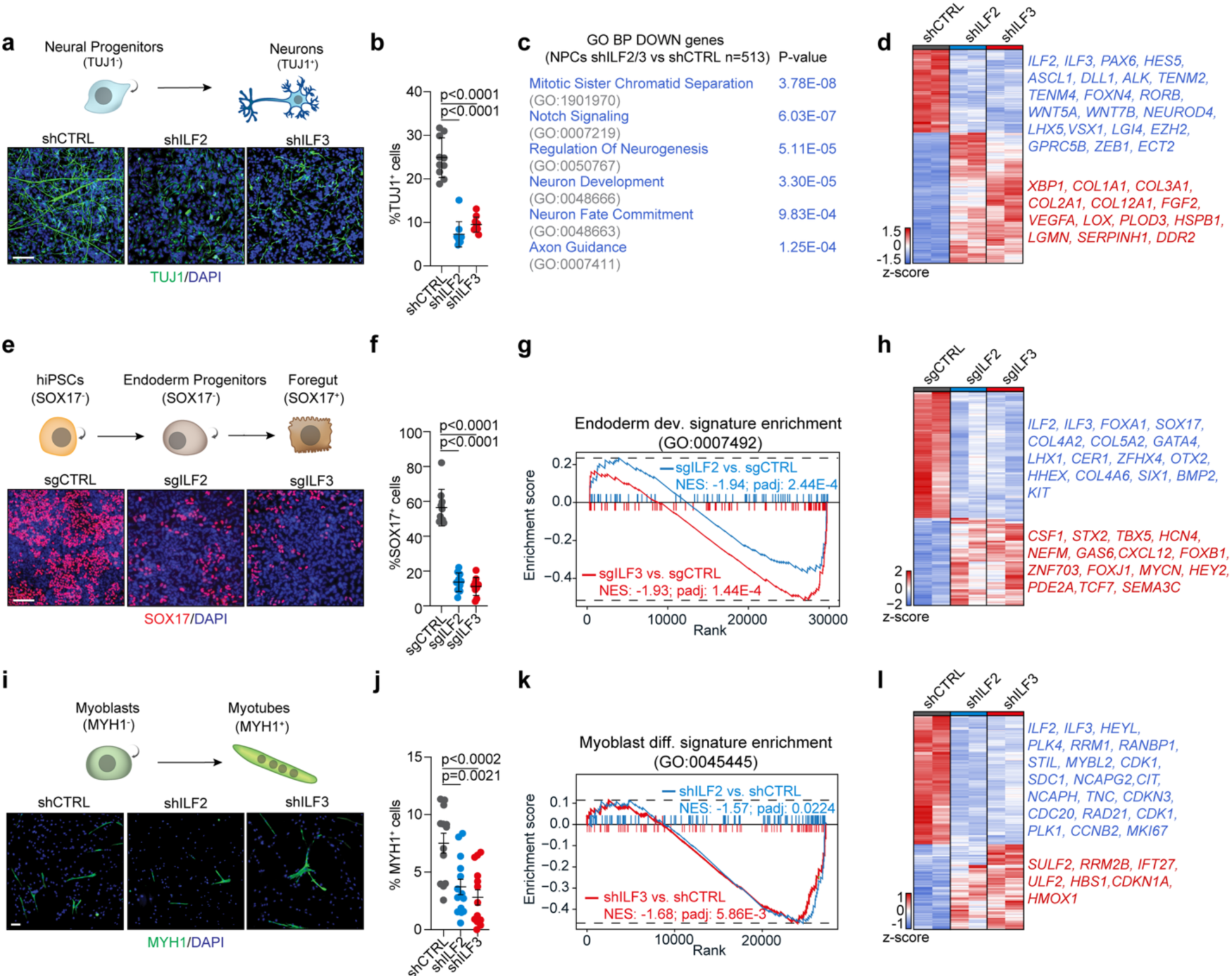
ILF2 and ILF3 are required for proper lineage specification across multiple human developmental contexts. (**a**) Immunofluorescence analysis of neuronal differentiation showing TUJ1 expression (green) and nuclear DAPI staining (blue) in control and *ILF2* or *ILF3*-depleted neural progenitor cells (NPCs). Scale bar, 100 µm. (**b**) Quantification of TUJ1-positive cells (n = 10 independent images per condition; P values determined by unpaired two-tailed Student’s t-test). (**c**) Gene Ontology enrichment analysis of Biological Processes (BP) of downregulated genes in *ILF2*- and *ILF3*-depleted neurons compared to controls (two-tailed Fisher’s exact test). (**d**) Differential gene expression analysis in neurons following *ILF2* or *ILF3* knockdown (n = 2 biological replicates; |fold change| > 1.5; P < 0.05, Wald test with Benjamini-Hochberg correction). Red and blue indicate up- and down-regulated genes, respectively. (**e**) Immunofluorescence analysis of endodermal differentiation showing SOX17 expression (red) and nuclear DAPI staining (blue) in control and *ILF2*/*3*-depleted foregut progenitors. Scale bar, 100 µm. (**f**) Quantification of SOX17-positive cells (n = 10 independent images per condition; P values determined by unpaired two-tailed Student’s t-test). (**g**) GSEA of endoderm-specific genes in *ILF2*- or *ILF3*-depleted foregut cells (shILF2: NES = -1.94, P = 2.44E-4; shILF3: NES = -1.93, P = 1.44E-4). (**h**) Differential gene expression analysis in foregut cells following *ILF2* or *ILF3* knockdown (n = 2 biological replicates; |fold change| > 1.5; P < 0.05, Wald test with Benjamini-Hochberg correction). Red and blue indicate up- and down-regulated genes, respectively. (**i**) Immunofluorescence analysis of myogenic differentiation showing MYH1 expression (green) and nuclear DAPI staining (blue) in control and *ILF2* or *ILF3*-depleted primary myoblasts. Scale bar, 100 µm. (**j**) Quantification of MYH1-positive cells (n = 8 independent images per condition; P values determined by unpaired two-tailed Student’s t-test). (**k**) GSEA of myoblast differentiation genes in *ILF2*- or *ILF3*-depleted myotubes (shILF2: NES = -1.57, P = 0.0224; shILF3: NES = -1.68, P = 5.86E-3). (**l**) Differential gene expression analysis in myotubes following *ILF2* or *ILF3* knockdown (n = 2 biological replicates; |fold change| > 1.5; P < 0.05, Wald test with Benjamini-Hochberg correction) Red and blue indicate up-and down-regulated genes, respectively.

To determine whether ILF2 and ILF3 regulate progenitor cell identity in lineages other than the ectoderm, we tested the effects of *ILF2* and *ILF3* suppression during differentiation of endoderm progenitors into foregut cells^74^ (**Fig. 2e and Extended Data Fig. 4d**). We found that *ILF2*- and *ILF3*-depleted endoderm cells failed to activate SOX17 in the majority of cells after 3 days of differentiation (**Fig. 2e,f**). Consistent with these findings, CRISPRi-mediated suppression of *ILF2* and *ILF3* impaired the activation of key foregut markers (**Fig. 2g and Extended Data Fig. 4e,f**), including *FOXA1, GATA4*, and *HHEX*^75, 76^ (**Fig. 2h and Extended Data Fig. 4g**). These results underscore the crucial roles of ILF2 and ILF3 in endoderm specification and differentiation.

Finally, to extend our observations to the mesoderm lineage, we investigated their roles in primary human myoblasts undergoing differentiation into myotubes. Primary human myoblasts were transduced with lentiviral vectors expressing shRNAs targeting *ILF2* or *ILF3*, or with control vectors, and assessed for the formation of MYH1^+^ myotubes after seven days (**Fig. 2i and Extended Data Fig. 4h**). Loss of *ILF2* or *ILF3* led to modest (∼2-fold) reduction in MYH1^+^ myotube formation (**Fig. 2i,j and Extended Data Fig. 4h**). Concordantly, GSEA analysis of RNA-seq data from control and *ILF2*- or *ILF3*-knockdown cells revealed partial downregulation of myoblast differentiation transcripts (**Fig. 2k,l and Extended Data Fig. 4i-k**). These findings suggest that ILF2 and ILF3 play a less prominent role in mesodermal differentiation compared to ectoderm and endoderm derivatives.

Together, these data indicate that ILF2 and ILF3 not only regulate the differentiation of PSCs but also facilitate adult progenitor cell differentiation, suggesting that ILF2 and ILF3 may act as critical gatekeepers of stem cell plasticity, particularly in the endoderm and ectoderm lineages.

### ILF2 and ILF3 modulate ADAR1-mediated A-to-I editing of transcripts encoding cell fate factors

DRBPs can interact with both DNA and RNA, with evidence suggesting initial recruitment to gene promoters followed by association with chromatin-associated transcripts^77^. To investigate the molecular mechanisms underlying ILF2 and ILF3 function in human stem cells, we first mapped the chromatin and RNA interaction landscapes of ILF3 in hPSCs. Using CUT&Tag^78^, we identified over 16,000 ILF3-chromatin-bound peaks (RPKM>0.5; top 1% of peaks by area under the curve (AUC))^79^, primarily localized to promoters, introns, and distal intergenic regions (**Fig. 3a and Extended Data Fig. 5a**). Transcription factor motif analysis revealed enrichment for cell fate regulators including Mediator, estrogen-related receptors, and POU homeodomain factors^76, 80–85^ (**Extended Data Fig. 5b**). These data support a model whereby ILF3 associates with chromatin at specific regulatory regions.

**Figure 3.**
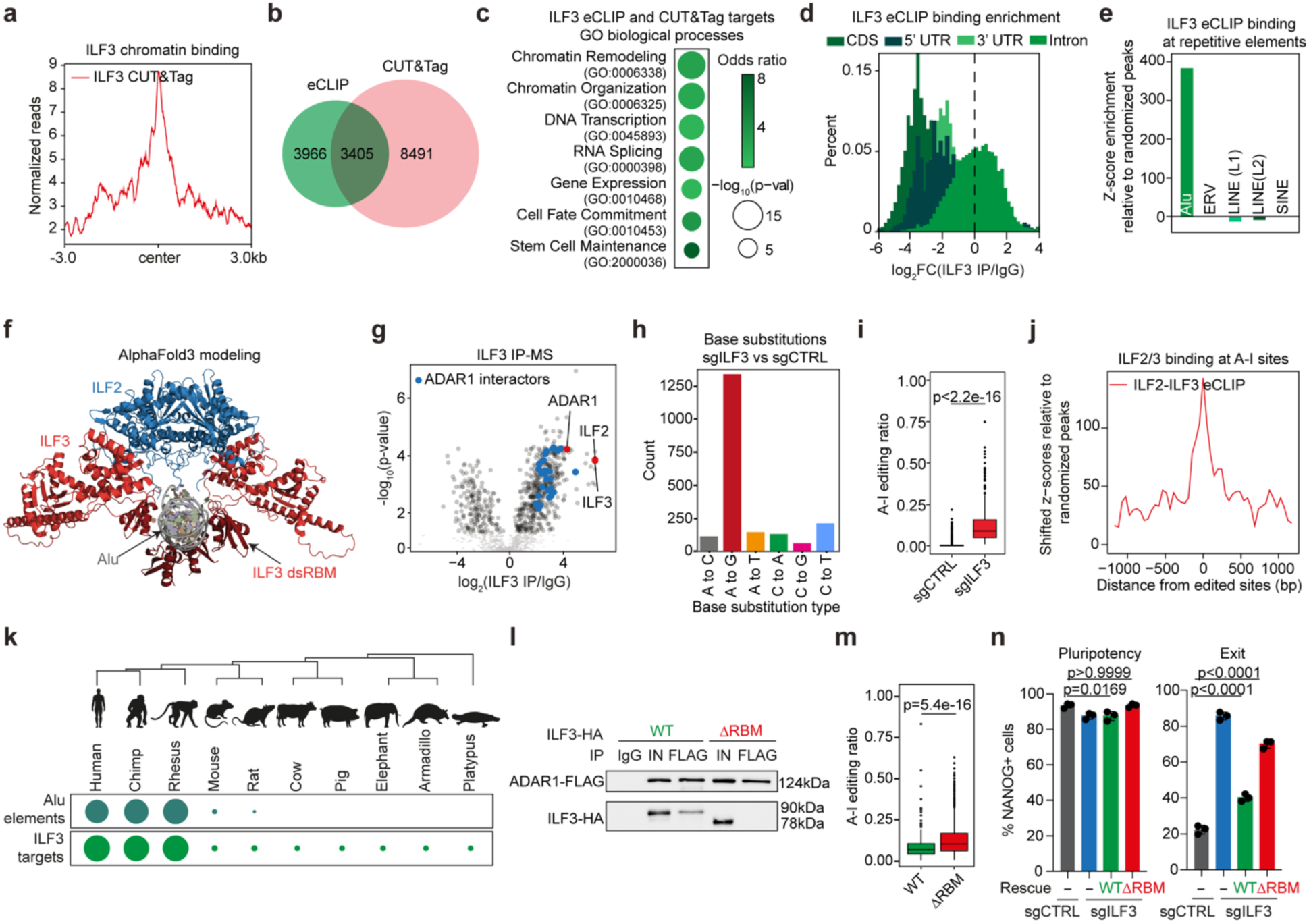
ILF3 regulates RNA editing through Alu element binding and ADAR1 interaction. (**a**) Genome-wide distribution of ILF3 binding sites determined by CUT&Tag analysis, showing normalized read density across gene bodies ±3 kb from transcriptional centers. (**b**) Overlap between ILF3 binding sites identified by CUT&Tag and eCLIP analyses (RPKM>0.5; top 1% of peaks by area under the curve (AUC). (**c**) Gene Ontology enrichment analysis of ILF3-bound regions identified by eCLIP-seq and CUT&Tag. (d) Distribution of ILF3 eCLIP signal intensity relative to size-matched input controls (log₂FC > 3; P < 0.001). (**e**) Enrichment analysis of repetitive element classes in ILF3 eCLIP peaks relative to a randomized peak distribution. (**f**) AlphaFold3-predicted structural model of the ILF2/3 complex bound to Alu RNA. (**g**) Proteomic analysis of ILF3 interactors identified by immunoprecipitation-mass spectrometry (n = 3 biological replicates). (**h**) Single nucleotide variants (SNVs) detected in ILF3-depleted versus control hPSCs. Base substitution types include counts for the reverse complement variant. (**i**) Comparison of editing frequencies at A-to-I edited sites in ILF3 knockdown or control knockdown hPSCs. (n = 2 biological replicates; P values determined by Wilcoxon rank-sum test). (**j**) Aggregate plot showing ILF2/3 eCLIP signal distribution centered around A-to-I-edited sites after ILF3 depletion. Z-scores calculated relative to a randomized peak distribution. (**k**) Phylogenetic tree – the diameter of each bubble is proportional to the percentage of each species’ respective genome that aligns to Alu elements (top row) or ILF3 eCLIP targets identified in hPSCs (bottom row). (**l**) Western blot analysis of interactions between FLAG-tagged ADAR1 and wild-type HA-ILF3 or RNA-binding mutant (HA-ILF3ΔRBM). (**m**) Editing frequencies in ILF3 wild-type versus ΔRBM rescue conditions (n = 2 biological replicates; P values determined by Wilcoxon rank-sum test). (**n**) Quantification of NANOG-positive cells under self-renewal and exit conditions following rescue with wild-type or ΔRBM mutant ILF3 (n = 3 biological replicates; P values determined by one-way ANOVA with Tukey’s multiple comparisons test).

Next, we used enhanced UV cross-linking and immunoprecipitation sequencing^86^ (eCLIP-seq) and identified over 7,000 ILF3-bound RNAs (*p<*0.05, log_2_FC>2.5), with strong correlation between replicates (**Fig. 3b and Extended Data Fig. 5c**). A subset of ILF3’s RNA targets overlap with its chromatin binding sites (∼46%), suggesting that ILF3 may engage with both chromatin and RNA at specific loci to support coordinated gene regulation (**Fig. 3b and Extended Data Fig. 5d,e**). Gene ontology analysis revealed enrichment for biological processes including “chromatin remodeling,” “cell fate commitment,” and “stem cell maintenance” (**Fig. 3c**). Given that the majority of ILF3-bound locations occurred in distal introns (**Fig. 3d and Extended Data Fig. 5f**), and considering the role of transposable elements in intron expansion^87–89^, we investigated whether ILF3 preferentially occupied specific transposable element subclasses. Strikingly, almost all ILF3 binding occurred within Alu elements, a family of primate-specific repetitive elements (**Fig. 3e and Extended Data Fig. 5g,h**). AlphaFold3 modeling^90^ predicted that the ILF2/3 complex binds Alu elements via double-stranded RNA-binding motifs (dsRBMs), adopting a structural configuration similar to that of other known Alu-binding proteins^91^ (**Fig. 3f**). This observation is consistent with recent evidence that the ILF2/3 complex can coat long segments of double-stranded RNAs containing Alus^92^. Analysis of ILF2 eCLIP-seq in two distinct hPSC lines, as well as ILF3 CUT&Tag data, reinforced these findings, showing preferential binding at Alu elements (**Extended Data Fig. 5i-m**). Collectively, our findings demonstrate that ILF2 and ILF3 target transcripts of cell fate regulatory proteins and may regulate their processing in a species-specific manner by interacting with Alu repetitive elements.

To define the mechanisms by which ILF2 and ILF3 control cell fate through Alu element binding, we determined the protein interactors of ILF2 and ILF3 in hPSCs using immunoprecipitation followed by mass spectrometry (IP-MS). Consistent with their known function as a heterodimer^52, 93, 94^, ILF2 and ILF3 co-precipitated during reciprocal pull-downs (**Fig. 3g and Extended Data Fig. 6a**). Among the strongest interactors, we identified ADAR1, an RNA deaminase that catalyzes adenosine-to-inosine (A-to-I) editing^95–98^ and is highly expressed in pluripotent cells (**Fig. 3g and Extended Data Fig. 6a,b**). ADAR1 shares key characteristics with ILF2 and ILF3: higher expression levels in humans relative to mice (**Extended Data Fig. 6b**), and preferential targeting of Alu elements^96–99^. We confirmed this interaction by performing IP of ILF3 followed by western blot, which specifically detected the nuclear p110 isoform of ADAR1^100^ (**Extended Data Fig. 6c**). We further validated this finding by FLAG-ADAR1 pull-down in cells co-expressing HA-tagged ILF3 in an independent hPSC line (**Extended Data Fig. 6d**). Together, these findings suggest that ILF2/3 may facilitate lineage specification in primates by modulating ADAR1-mediated RNA editing at Alu elements.

To determine whether ILF2 and ILF3 influence ADAR1 activity and RNA editing, we analyzed editing levels following *ILF3* depletion in hPSCs and adult progenitors. Since silencing *ILF3* is sufficient to deplete both proteins (**Fig. 1b**) and 93% of ILF2 targets are also bound by ILF3 (**Extended Data Fig. 6e**), we utilized ILF3 depletion as a proxy to study the effects of disrupting the ILF2/3 complex on A-to-I editing. RNA sequencing and single nucleotide variant analysis stratified by base changes revealed significant increases in A-to-I editing following *ILF3* depletion (**Fig. 3h,i**). Importantly, ILF2 and ILF3 binding was enriched at sites showing increased RNA editing, suggesting a direct role for the ILF2/3 complex in suppressing this process (**Fig. 3j**). To further investigate how ILF2/3 affects ADAR1 function, we performed ADAR1 RNA immunoprecipitation followed by sequencing (RIP-seq) after *ILF3* knockdown. Our analysis confirmed preferential binding of ADAR1 at intronic regions^101^ (**Extended Data Fig. 6f**), mirroring the binding pattern of ILF2/3. Notably, upon ILF3 depletion, we observed enhanced ADAR1 binding to ILF2/3 targets, including transcripts that undergo increased editing in ILF3-deficient cells (**Extended Data Fig. 6g,h**). These findings suggest that the ILF2/3 complex functions as a negative regulator of ADAR1-mediated RNA editing at specific target sites.

To extend these observations beyond hPSCs and determine conservation of this regulatory mechanism across different lineages, we analyzed RNA editing levels following *ILF3* depletion in adult progenitors. *ILF3* suppression significantly increased A-to-I editing in neural and endoderm progenitors, with more modest changes in human myoblasts (**Extended Data Fig. 6i**), corroborating our phenotypic observations across the three lineages and highlighting a lineage-specific role for ILF3 in suppressing RNA editing.

Finally, our data reveal that genomic sequences corresponding to ILF3-bound RNAs identified in hPSCs are significantly more conserved in primates compared to other species, such as mice (**Fig. 3k**). This conservation pattern suggests that primates may have co-opted the ILF2/3 complex to regulate ADAR1-mediated RNA editing at Alu elements. We thus hypothesized that if RNA misediting occurs primarily at primate-specific sequences and is functionally linked to ILF2 and ILF3 in cell fate regulation, then depleting these factors in mouse ESCs—where no substantial phenotype is observed following ILF2/3 loss—should not lead to increased RNA editing. To test this, we performed RNA sequencing and A-to-I editing analysis at ILF3-bound regions^102^ following *Ilf2* and *Ilf3* knockdown or knockout in mouse pluripotent cells^102^. In mouse ESCs, the ILF2/3 complex did not preferentially bind B1 repetitive elements, which are structurally similar to human Alu sequences^103^ (**Extended Data Fig. 6j**). Strikingly, and in contrast to our observations in hPSCs, knockdown of *Ilf2* or *Ilf3* in mouse ESCs did not result in dramatic changes in editing levels at ILF3-binding sites (**Extended Data Fig. 6k**).

Together, these findings establish ILF2 and ILF3 as key interactors of ADAR1 and modulators of A-to-I RNA editing at Alu elements. This primate-specific mechanism may explain the species-specific requirement for ILF2/3 in human, but not mouse, stem cell fate determination, providing a molecular basis for evolutionary divergence in developmental regulation.

### The double-stranded RNA-binding motifs of ILF3 are essential for inhibiting RNA-editing and cell fate transitions

ILF3 and ADAR1 both contain double-stranded RNA-binding motifs (dsRBMs), which enable binding to Alu elements that form double-stranded RNAs during transcription^95, 104–108^. To test whether the dsRBM domains of ILF3 are indeed required for its interaction with ADAR1, we generated HA-tagged constructs expressing either ILF3 wild-type (ILF3WT) or an ILF3 variant lacking both dsRBMs^94, 109–111^ (ILF3ΔRBM, **Extended Data Fig. 6l**). These constructs were co-transfected into hPSCs alongside FLAG-tagged ADAR1p110, followed by FLAG-ADAR1 immunoprecipitation. While ILF3WT robustly co-immunoprecipitated with ADAR1, the interaction was completely abolished in ILF3ΔRBM-expressing cells (**Fig. 3l**). To determine whether the interaction between ILF2/3 and ADAR1 depends on DNA or RNA, we treated hPSC lysates with benzonase^112^ to degrade nucleic acids. Notably, this treatment did not disrupt the binding between ILF2/3 and ADAR1 (**Extended Data Fig. 6m**). These results indicate that the dsRBM domains of ILF3 are indispensable for its interaction with ADAR1 and that maintenance of this interaction occurs independently of RNA.

If disrupting the ILF3-ADAR1 interaction underlies the phenotypic effects observed in *ILF2*- and *ILF3*-depleted cells, we reasoned that the dsRBM domains of ILF3 would be necessary for the proper RNA editing and differentiation of embryonic stem cells. To test this possibility, we performed rescue experiments in our CRISPRi ILF3 hPSCs with either ILF3WT or ILF3ΔRBM (**Extended Data Fig. 6n,o**). RNA-seq analysis showed that A-to-I editing was significantly increased in the ILF3ΔRBM cells compared to their wild-type counterparts at regions directly bound by ILF2/3 (**Fig. 3m and Extended Data Fig. 6p**). These editing changes recapitulated the misediting phenotype observed following ILF3 depletion, underscoring the critical role of ILF3 in suppressing RNA editing through its dsRBM domains. To evaluate the functional impact of the dsRBM domains on cell fate, we induced differentiation of ILF3 CRISPRi cells rescued with either ILF3WT or ILF3ΔRBM and monitored NANOG expression. While ILF3WT-rescued cells regained differentiation capacity, as evidenced by the downregulation of NANOG, ILF3ΔRBM-rescued cells largely failed to downregulate NANOG, indicating impaired differentiation (**Fig. 3n**).

In summary, our findings demonstrate that the dsRBM domains of ILF3 are essential for its interaction with ADAR1 and for repressing A-to-I RNA editing. Furthermore, these domains are critical for enabling proper cell fate transitions in hPSCs.

### ILF2 and ILF3 depletion causes mis-splicing of transcripts encoding fate-instructive proteins

Increased A-to-I editing at Alu elements within intronic regions has been reported to induce the inclusion of these regions into mature transcripts, a process referred to as exonization^30, 104, 113–118^. Building on our observation that *ILF2* and *ILF3* depletion induces hyper-editing at Alu elements, we hypothesized that ILF3 influences alternative splicing of transcripts encoding key cell fate regulators. To test this, we established an ILF3 degron system in hPSCs to study early, direct splicing changes caused by ILF3 loss. Using CRISPR/Cas9, we introduced an FKBP12^F36V^-HA-P2A-mCherry cassette into the *ILF3* locus. Treatment with dTAG^V^-1^119^ resulted in rapid ILF3 protein degradation and concomitant loss of ILF2 protein within 24 hours, thus efficiently depleting the ILF2/3 complex without affecting *ILF2* or *ILF3* mRNA levels (**Fig. 4a and Extended Data Fig. 7a**).

**Figure 4.**
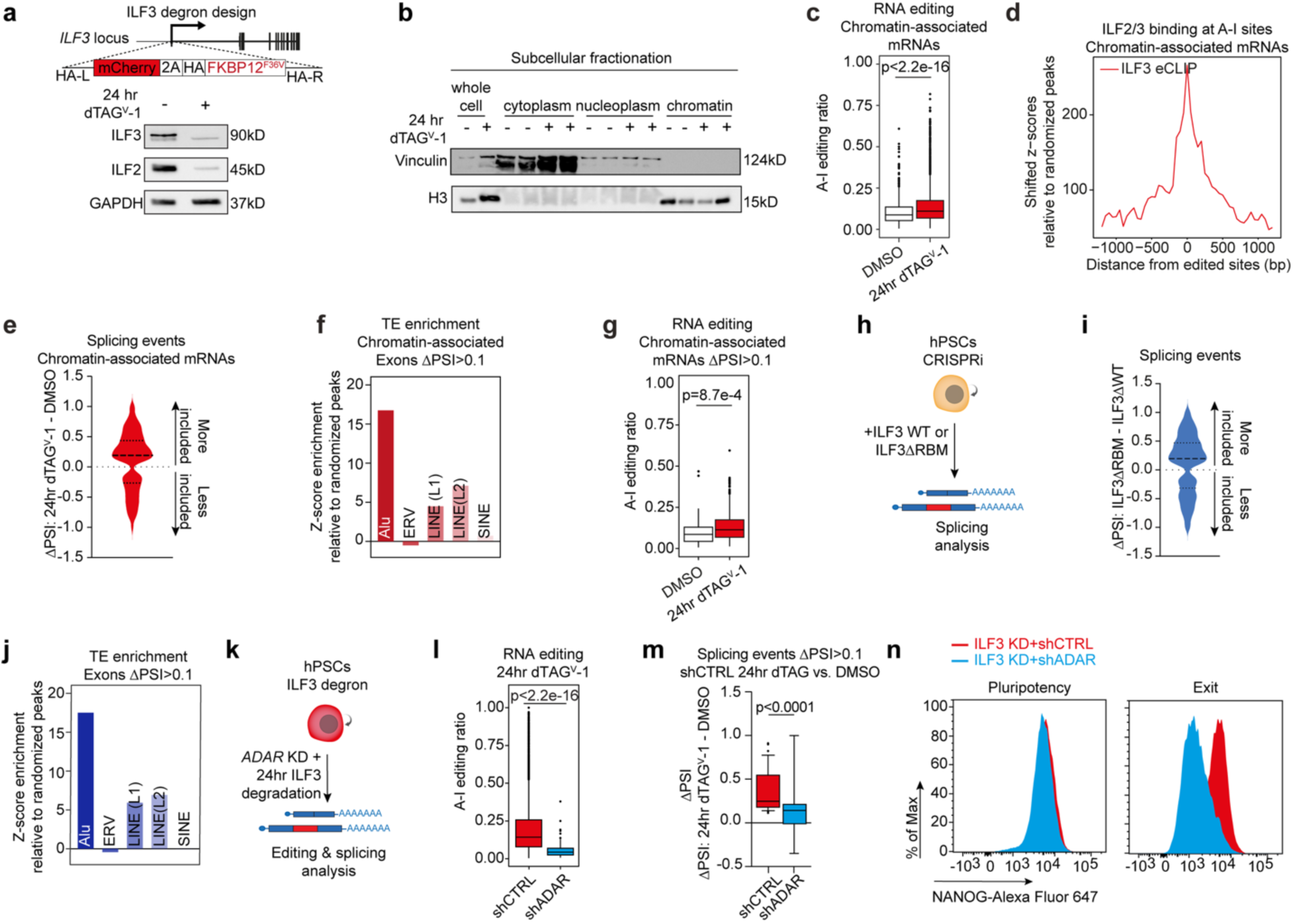
Acute ILF3 degradation triggers widespread splicing defects in hPSCs. (**a**) Generation of ILF3-FKBP12^F36V^ hPSCs using CRISPR-Cas9-mediated knock-in (top) and western blot showing ILF3 and ILF2 protein levels after 24 hours of dTAG^V^-1 treatment (bottom). (**b**) Western blot analysis of fractionated whole cell and subcellular lysates. (**c**) Editing frequencies in chromatin-associated mRNAs after 24 hours of ILF3 degradation (n = 2 biological replicates; P values determined by Wilcoxon rank-sum test). (**d**) Aggregate plot showing ILF3 eCLIP signal distribution centered around A-to-I-edited sites in chromatin-associated mRNAs after 24 hours of ILF3 degradation. Z-scores calculated relative to a randomized peak distribution. (**e**) Alternative splicing events in chromatin-associated mRNAs 24 hours after ILF3 degradation (|ΔPSI| > 0.1, FDR < 0.05). (**f**) Enrichment of repetitive elements in cassette exons more included in chromatin-associated mRNAs after ILF3 degradation (ΔPSI > 0.1, FDR < 0.05) relative to a randomized peak distribution. (**g**) Editing frequencies in transcripts showing increased exon inclusion after ILF3 degradation in chromatin-associated mRNAs (ΔPSI > 0.1, FDR < 0.05; n = 2 biological replicates; P values determined by Wilcoxon rank-sum test). (**h**) Experimental design for splicing analysis in CRISPRi ILF3 hPSCs after rescue with ILF3 WT or ILF3ΔRBM. (**i**) Alternative splicing comparison between ILF3ΔRBM and wild-type rescue in *ILF2*/*3*-depleted cells (|ΔPSI| > 0.1, FDR < 0.05). (**j**) Enrichment of repetitive elements in cassette exons more included in ILF3ΔRBM versus wild-type rescue (ΔPSI > 0.1, FDR < 0.05) relative to a randomized peak distribution. (**k**) Experimental design for editing and splicing analysis after ILF3 degradation and *ADAR* knockdown. (**l**) Editing frequencies at A-to-I sites observed after 24 hours of ILF3 degradation in *ADAR* knockdown hPSCs, relative to events observed in control knockdown cells. (n = 2 biological replicates; P values determined by Wilcoxon rank-sum test). (**m**) Alternative splicing ratios 24 hours after ILF3 degradation in *ADAR* knockdown hPSCs, relative to inclusion events observed in control knockdown cells (ΔPSI > 0.1, FDR < 0.05; P values determined by Wilcoxon rank-sum test). (**n**) Representative flow cytometric analysis of NANOG-positive cells under self-renewal and exit conditions following *ILF3* silencing and rescue with control or *ADAR* knockdown.

RNA editing by the nuclear isoform ADAR1 occurs co-transcriptionally^120^, raising the question of whether ILF3 affects this process during early stages of transcriptional processing. To address this, we performed subcellular fractionation to isolate chromatin-associated transcripts from ILF3 degron hPSCs, followed by analysis of RNA editing and splicing changes^121^ (**Fig. 4b**). If ILF3 acts on newly transcribed RNAs, we would expect to observe editing changes in chromatin-bound transcripts shortly after ILF3 degradation. Indeed, chromatin-associated transcripts showed significant increases in A-to-I editing at ILF2/3-bound RNAs within 24 hours (**Fig. 4c,d**). Notably, this rapid increase in RNA editing coincided with significant changes in alternative splicing, characterized by greater inclusion of exons that were enriched for Alu elements (**Fig. 4e-g**). Analysis of RNA extracted from whole cell lysates after ILF2/3 degradation displayed similar editing and splicing patterns (**Extended Data Fig. 7b-j**). These findings demonstrate that the ILF2/3 complex controls ADAR1 activity in the nucleus, such that ILF3 degradation induces hyperediting and exonization of target transcripts shortly after transcription.

We next extended our analysis to adult progenitor cells, including NPCs, myoblasts, endoderm progenitors, in addition to mouse ESCs. Similar to hPSCs, ILF3 depletion enhanced exonization in NPCs and endoderm progenitors, with a more modest effect in myoblasts (**Extended Data Fig. 7k**), consistent with observed RNA editing changes and phenotypes observed upon *ILF2* and *ILF3* suppression (**Fig. 2i,j and Extended Data Fig. 6i**). As expected, *Ilf2* and *Ilf3* depletion in mouse ESCs did not increase intron inclusion, consistent with the lack of hyper-editing *in Ilf2* and *Ilf3*- depleted cells (**Extended Data Figs. 6k and 7l**).

To corroborate these findings, we used a pipeline designed to detect splice junctions between exons and transposable elements (”JETs”^29, 122^) in hPSCs. Following ILF3 depletion, we identified 93 differentially spliced JETs, most of which were more abundant in ILF3-depleted cells (**Extended Data Fig. 7m**). These JET-containing transcripts were enriched for Alu elements and A-to-I editing (**Extended Data Fig. 7n,o**), further linking ILF3 loss to transposable element-driven splicing dysregulation.

Next, we examined whether ADAR1 or its interaction with ILF3 was required for the observed changes in transcript splicing. To this end, we compared splicing patterns in CRISPRi hPSCs expressing either ILF3ΔRBM, which is unable to interact with ADAR1 (**Fig. 3l**), or wild-type ILF3. Our analysis revealed significantly increased inclusion of Alu-containing introns in ILF3ΔRBM-rescued cells compared to the wild-type control (**Fig. 4h-j**), indicating that the ILF3-ADAR1 interaction is important for proper editing and splicing of ILF3-target transcripts. To further establish the cooperative role of ILF3 and ADAR1 in RNA splicing, we analyzed splicing patterns in ILF3 degron cells (24 hours post-depletion) infected with lentiviral vectors expressing *ADAR*-targeting shRNAs or a control vector (**Fig. 4k and Extended Data Fig. 7p**). *ADAR* depletion markedly reduced RNA editing events induced by ILF3 degradation (**Fig. 4l**). Furthermore, *ADAR* knockdown partially rescued alternative splicing patterns and impeded the formation of JETs normally induced by ILF3 degradation (**Fig. 4m and Extended Data Fig. 7q,r**). These results suggest that ADAR1-mediated hyperediting at Alu elements drives the exonization observed following ILF3 depletion. We reasoned that if *ADAR* depletion mitigates splicing defects caused by ILF3 degradation, it might also rescue the associated differentiation defect. Indeed, knockdown of *ADAR* in ILF3-depleted hPSCs restored NANOG downregulation during exit from pluripotency (**Fig. 4n**), supporting a role for the ILF3-ADAR1 axis in safeguarding proper cell fate transitions.

Together, these results demonstrate that ILF3 regulates A-to-I editing to prevent aberrant exonization of transposable elements in transcripts. This mechanism is essential in human, but not mouse, PSCs as well as in human adult progenitor cells, underscoring a species-specific role for ILF3 in splicing regulation.

### Mis-spliced transcripts formed after ILF2 and ILF3 degradation undergo decay

Alu exonization can introduce premature termination codons, leading to transcript degradation via the nonsense-mediated decay (NMD) pathway^30, 123^. We hypothesized that transcripts undergoing increased exonization following ILF3 degradation might be preferentially targeted for decay through this mechanism. Consistent with this hypothesis, while ILF3 depletion resulted in comparable numbers of up- and down-regulated transcripts in hPSCs (**Extended Data Fig. 3a**), transcripts exhibiting Alu exonization were significantly downregulated after 24 hours of ILF3 degradation, a trend maintained at 96 hours (**Fig. 5a and Extended Data Fig. 7s**). Furthermore, ILF3 depletion led to a marked increase in inclusion of premature termination codons specifically within ILF3 target genes (**Fig. 5b**), suggesting that exonized transcripts in ILF3-depleted cells might undergo degradation via NMD.

**Figure 5.**
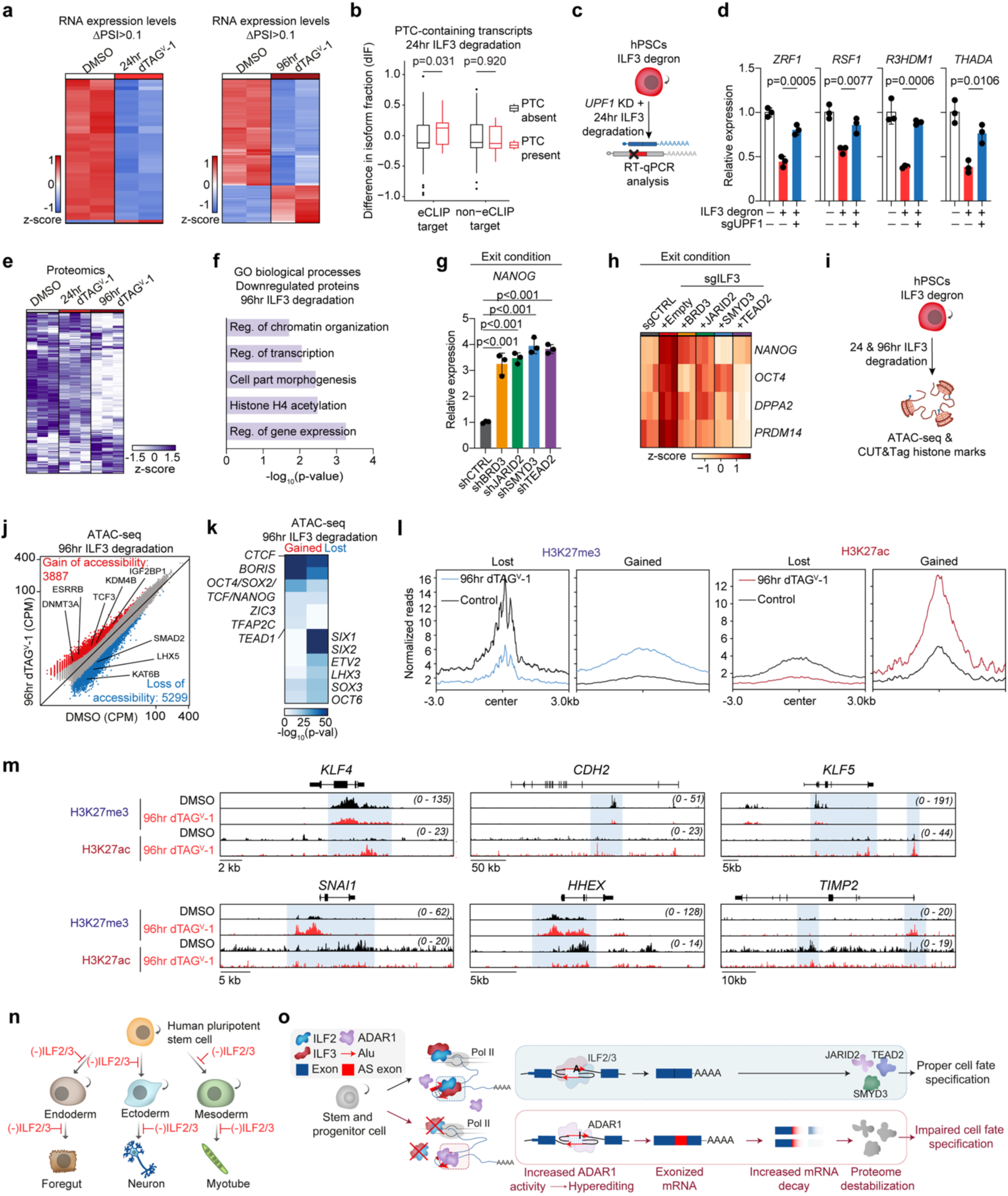
ILF3 degradation rewires the chromatin and proteome landscapes in hPSCs. (**a**) RNA-seq expression analysis of mis-spliced transcripts after 24 (left) and 96 hours (right) of ILF3 degradation (FDR < 0.05; ΔPSI > 0.1; |FC| > 1.5, P < 0.05). (**b**) IsoformSwitch analysis of PTC-containing vs. non-PTC-containing transcript isoforms after 24 hours of ILF3 degradation, divided by ILF3 eCLIP targets (left) and non-targets (right). P values determined by Kolmogorov-Smirnoff test. (**c**) Experimental design for transcript analysis after ILF3 degradation and *UPF1* knockdown. (**d**) RT-qPCR analysis of mis-spliced transcripts with and without *UPF1* knockdown (n = 3 biological replicates; mean ± s.d.; P values determined by one-way ANOVA with Tukey’s multiple comparisons test). (**e**) Proteomic analysis of mis-spliced transcripts after 24 and 96 hours of ILF3 degradation (FDR < 0.05; ΔPSI > 0.1; P < 0.05). (**f**) GO analysis of downregulated proteins associated with mis-spliced transcripts. (**g**) RT-qPCR analysis of *NANOG* expression after knockdown of chromatin regulators in hPSCs under exit conditions induced by MAPK pathway inhibition (n = 3 biological replicates; mean ± s.d.; P values determined by one-way ANOVA with Dunnett’s multiple comparisons test). (**h**) RT-qPCR analysis of pluripotency gene expression after control or *ILF3* knockdown with and without rescue by indicated genes in hPSCs under exit conditions (n = 3 biological replicates). (**i**) Experimental design for chromatin and histone mark analysis after ILF3 degradation. (**j**) ATAC-seq analysis after 96 hours of ILF3 degradation (n = 2 biological replicates; red: increased accessibility, blue: decreased accessibility; |FC| > 1.5, P < 0.05). (**k**) HOMER motif analysis at differential chromatin accessibility regions (n = 2 biological replicates; |FC| > 1.5; P < 0.05). (**l**) Aggregate histone modification profiles after 96 hours of ILF3 degradation (n = 3 biological replicates) versus control. (**m**) Representative CUT&Tag tracks showing changes in histone mark deposition at pluripotency (left) and differentiation (right) genes. (**n**, **o**) Proposed mechanistic model (AS = Alternative spliced).

To directly assess whether NMD mediates the decay of these exonized transcripts, we inhibited *UPF1*, a core NMD factor essential for degrading mis-spliced RNAs^30, 124, 125^. We engineered ILF3 degron hPSCs to incorporate a CRISPRi system for inducible and efficient *UPF1* depletion upon doxycycline treatment (**Fig. 5c and Extended Data Fig. 7t**). We reasoned that if NMD drives the degradation of ILF3-targeted RNAs that undergo Alu exonization, *UPF1* silencing should restore their expression. Indeed, *UPF1* depletion significantly rescued the expression of transcripts downregulated following ILF3 degradation in dTAG^V^-1-treated cells (**Fig. 5d**).

Together, these results demonstrate that transcripts mis-spliced due to ILF3 loss are degraded via UPF1-mediated decay, highlighting a critical role for ILF2 and ILF3 in maintaining transcript integrity and stability.

### ILF2 and ILF3 ensure proteome fidelity and regulate chromatin architecture in hPSCs

To determine whether degradation of mis-spliced transcripts following ILF3 depletion impacts protein levels, we performed proteomic analysis in ILF3 degron hPSCs at 0, 24, and 96 hours post-dTAG^V^-1 treatment (**Extended Data Fig. 8a**). By day 4, we observed that exonized transcripts showing detectable changes at the protein level were predominantly downregulated (**Fig. 5e**), consistent with our RNA expression data (**Fig. 5a and Extended Data Fig. 7s**). Gene ontology analysis revealed that the downregulated proteins were enriched for categories related to chromatin regulation and transcription (**Fig. 5f**). For example, downregulated proteins included chromatin-associated factors such as BRD3, JARID2, SMYD3, and TEAD2. These results raised the possibility that degradation of mis-spliced chromatin-associated factors contributes to the impaired cell fate transitions observed following ILF2/3 depletion. Consistent with this model, suppression of *BRD3*, *JARID2*, *SMYD3*, and *TEAD2* in hPSCs blocked their exit from pluripotency, mimicking the ILF2/3 phenotype (**Fig. 5g and Extended Data Fig. 8b**). Moreover, forced expression of correctly spliced chromatin factors after ILF2/3 silencing partially rescued the capacity of cells to downregulate pluripotency-associated transcripts (**Fig. 5h**). Together, these findings demonstrate that ILF3 suppression drives exonization, leading to reduced RNA and protein levels of key chromatin factors, ultimately impairing cell fate transitions.

The observation that ILF3 modulates the editing and splicing of key chromatin regulators raised the question of whether ILF3 degradation alters chromatin organization in pluripotent cells. To test this, we performed ATAC-seq analysis in hPSCs following 24 and 96 hours of ILF3 degradation (**Fig. 5i**). ILF3-depleted hPSCs exhibited a global decrease in accessible chromatin regions compared to controls (2817 sites with increased accessibility and 3953 sites with decreased accessibility after 24 hours; 3887 sites with increased accessibility and 5299 sites with decreased accessibility after 96 hours) (**Fig. 5j and Extended Data Fig. 8c**). Regions retaining accessibility were enriched for binding motifs of key pluripotency factors, including OCT4, SOX2, TCF, NANOG, and TFAP2C^12, 126, 127^ (**Fig. 5k**). In contrast, regions losing accessibility were associated with differentiation-related processes and transcription factor motifs (**Fig. 5k and Extended Data Fig. 8d**).

To further investigate how ILF3 depletion reshapes the chromatin landscape in hPSCs, we assessed the genome-wide distribution of histone marks linked to transcriptional activation and repression during cell fate transitions using CUT&Tag. We observed significant redistribution of the enhancer-associated histone mark H3K27ac and the Polycomb-associated histone mark H3K27me3, alongside more modest redistributions of H3K36me3 and H3K4me3 (**Fig. 5l and Extended Data Fig. 8e,f**). These changes included increased H3K27ac and concomitant decrease in H3K27me3 at pluripotency genes such as *CDH2* and *KLF4*^128, 129^, while the inverse pattern occurred at lineage-specifying genes such as *SNAI1* and *HHEX*^130, 131^ (**Fig. 5m**).

Collectively, these findings demonstrate that ILF3 loss initiates a sequential cascade whereby RNA misediting at Alu elements leads to transcript mis-splicing and degradation, destabilizing chromatin regulators and reshaping the epigenetic landscape at fate-instructive loci. These events compromise stem cell potency, revealing a mechanistic basis for the essential role of ILF2/3 in primate development.

## DISCUSSION

Our results reveal an evolutionary molecular safeguard that enables species-specific cell fate regulation in primates and highlight a role for RNA editing regulation during early embryonic cell fate transitions. We uncovered ILF2 and ILF3 as critical regulators of human cell identity (**Fig. 5n**). These proteins are essential for stem and progenitor cell differentiation and are required for primate gastrulation but dispensable in mice. Mechanistically, the ILF2/3 complex binds introns of fate-instructive transcripts, preventing ADAR1-directed RNA editing at primate-specific repetitive sequences. Loss of these proteins triggers transcript mis-splicing and decay, driving widespread alterations in protein expression and chromatin architecture (**Fig. 5o**).

Our findings reveal a striking example of evolutionary co-option, wherein the ILF2/3 complex has acquired a specialized function in regulating ADAR1-mediated RNA editing at Alu elements in primates, with direct consequences for pluripotent stem cell fate determination. This mechanism represents a compelling case of molecular repurposing during evolution, similar to the repurposing of crystallins from stress proteins to lens proteins^132^, TRIM5 from a general antiviral factor to an HIV-specific restriction factor in Old World monkeys^133^, and CTCF’s co-option for imprinting control in placental mammals^134^. While ILF2 and ILF3 maintain their ancestral roles in RNA metabolism in both mice and primates^135–137^, our data suggest primates have uniquely co-opted this complex to regulate A-to-I editing at Alu elements, possibly to manage the increased complexity of post-transcriptional regulation necessitated by repetitive element expansion in primate genomes^138–140^.

The evolutionary expansion of Alu elements in primates created a unique regulatory challenge^141^. While murine B1 repeats structurally resemble Alu elements, humans exhibit a 12-fold increase in long dsRNA formation, potentially explaining the 10-fold higher ADAR activity compared to mice^142^. These distinctions may underlie the divergent roles of ILF2 and ILF3 in mice versus humans^102^. Furthermore, the unique functions of ILF2 and ILF3 in regulating Alu elements suggest substantial evolutionary pressure to constrain the impact of Alu repeats on the human transcriptome^118, 143, 144^. This evolutionary adaptation may explain how human cells have co-opted ILF2 and ILF3 to safeguard transcriptome integrity and regulate cellular identity.

The molecular basis for the functional adaptation of ILF2 and ILF3 likely involves subtle evolutionary changes rather than major structural alterations. While the RNA-binding domains of these proteins are highly conserved, the functional adaptation may involve changes in protein-protein interactions, alterations in cellular localization, or context-dependent regulatory mechanisms. The heterodimeric nature of ILF2/3—a relatively recent evolutionary development that precludes homodimer formation^93, 145, 146^—enables functional diversification, allows independent regulation of each subunit’s expression across different contexts, and facilitates binding to asymmetric RNA structures formed by Alu elements^92^, highlighting how evolutionary tinkering with existing molecular machinery has generated novel regulatory mechanisms controlling cell fate decisions in primates.

We show that the inhibitory function of ILF3 on ADAR1-mediated editing in hPSCs relies on its double-stranded RNA-binding motifs (dsRBMs). Recent work demonstrated that the ILF2/3 complex can bind extensively along its targets, effectively “coating” long dsRNA sequences and potentially blocking the binding of other proteins^92^. One possibility is that ILF2/3 binding to dsRNAs prevents the editase domain of ADAR1 from accessing its targets by promoting structural alterations such as quadruplex formation. However, our results suggest that ILF3 and ADAR1 may also interact without DNA or RNA (**Extended Data Fig. 6m**)^100^. An additional possibility is that ILF2 and ILF3 might competitively inhibit ADAR1 by sequestering it away from RNA targets, thereby preventing editing. These inhibitory strategies bear similarities to splicing regulation by factors like heterogeneous nuclear ribonucleoproteins (hnRNPs), which can block factor recognition, inhibit assembly, or alter RNA structure to modulate splicing^147–152^.

An intriguing observation is that *ILF2* and *ILF3* depletion has a more pronounced effect on RNA editing in endoderm and ectoderm lineages compared to mesoderm. Mesodermal tissues may require tighter regulation of RNA editing and rely on alternative complexes to suppress ADAR1 activity, consistent with RNA-sequencing data showing that muscle tissue has among the lowest *ADAR* expression and A-to-I editing in mammals^153–155^. This lineage-specific requirement may reflect evolutionary constraints on RNA editing regulation in different developmental programs.

Proteomic analysis following ILF2 and ILF3 depletion revealed enrichment of chromatin-associated factors. Consistent with this finding, we observed significant changes in chromatin accessibility and histone mark distribution following *ILF2* and *ILF3* suppression, particularly in H3K27me3 and H3K27ac marks. While the molecular mechanisms by which ILF2 and ILF3 depletion affects these histone modifications remain incompletely understood, several factors likely contribute to the observed changes. For instance, JARID2, a component of the Polycomb complex important for regulating H3K27me3 deposition in pluripotent stem cells^156–158^ and SMYD3, known to catalyze trimethylation of H3K4^159, 160^, partially contribute to the effects observed after ILF2/3 depletion.

Dysregulated RNA editing has been implicated in a range of diseases, including cancer, autoimmune disorders, diabetes, and neurological conditions. Given the association of ILF2 and ILF3 expression with myeloma^51, 161^ and their involvement in autoimmune and neurodevelopmental disorders^46, 162, 163^, we propose that their roles in these pathologies may be linked to their function in modulating RNA editing. It will be critical to determine whether the mechanism we uncovered can be leveraged to better understand disease progression or inform therapeutic interventions.

In summary, our work identifies a molecular node that appears to have evolved to safeguard the integrity of the human transcriptome and drive cell fate decisions. This mechanism is crucial for maintaining proteome fidelity and preserving chromatin architecture, thereby enabling primate lineage specification. More broadly, our findings suggest that the co-option of DRBPs may represent a key evolutionary strategy that enabled the increased developmental complexity observed in primates. Future investigations into additional species-specific mechanisms will provide deeper insights into how these pathways coordinate cell fate and how their dysregulation contributes to the pathogenesis of complex diseases, including congenital malformations and cancer.

## Acknowledgements

We thank all members of the Di Stefano lab for helpful scientific discussions. Chimp iPSCs were a generous gift from Dr. Yoav Gilad. B.D.S. is a Cancer Prevention and Research Institute of Texas (CPRIT) Scholar in Cancer Research. B.D.S. is additionally supported by the CPRIT Recruitment of First-Time, Tenure-Track Faculty Member Award RR200079, the American Society of Hematology Scholar Award, the American Cancer Society award RSG-25-1510146-01-RMC, the B+ Foundation, the Worldwide Cancer Research Foundation, NIH awards (NIGMS MIRA R35GM147126, NIAID R21AI193649, and NCI R01CA291649). E.J.P. is supported by the Bert W. O’Malley, M.D. Scholar in Medical Research award funded by the Diana Helis Henry Medical Research Foundation and the Adrienne Helis Mavin Medical Research Foundation. E.J.P. is also supported by NIH award NICHD F30HD114315. C.M.S. is supported by the ASH Graduate Hematology Award. The authors acknowledge support by the state of Baden-Württemberg through bwHPC and the German Research Foundation (DFG) through grant INST 35/1597-1 FUGG. J.J.C. is supported by the NIH grant P41GM108538. A.W. is funded by the NIH grant R01HD112488 and the NSF grant MCB-2135296. This project was supported by the Cytometry and Cell Sorting Core at Baylor College of Medicine with funding from the CPRIT Core Facility Support Award (CPRIT-RP180672), the NIH (CA125123 and RR024574) and the assistance of Joel M. Sederstrom.

## Contributions

E.J.P. and B.D.S. conceived the study and wrote the manuscript; Y.C. performed the bioinformatics analysis; E.J.P. and F.L. conducted molecular biology and tissue culture experiments; V.L.S. and M. H. performed the ILF3 and histone mark CUT&Tag analysis; H. W. and J. W. the three-dimensional human peri-gastruloid models and analysis; N.L.G. the co-IP experiments; C.M.S. the IP-MS experiments; P.P. the endoderm differentiation and mouse gastruloid experiments; J.B. protein structural analyses; L.C. and J.C. conducted the proteomics; K.C. and H.C.H. RNA editing analyses; M.C.G. and A.W. produced the two-dimensional human gastruloid models. L.M. performed analysis of chromatin binding; E.V.N. assisted with performance and analysis of the ILF2 and ILF3 eCLIP experiments.

## Ethics declarations

E.V.N. is co-founder, member of the Board of Directors, on the SAB, equity holder and paid consultant for Eclipse BioInnovations, on the SAB of RNAConnect and is inventor of intellectual property owned by University of California San Diego. The interests of E.V.N. have been reviewed and approved by Baylor College of Medicine in accordance with its conflict of interest policies. J.J.C. is a consultant for Thermo Fisher Scientific, 908 Devices and Seer. The remaining authors declare no competing interests.

## METHODS

### Ethics statement

Human pluripotent stem cell experiments were performed at Baylor College of Medicine, Rice University, and the UT Southwestern Medical Center and followed the 2021 Guidelines for Stem Cell Research and Clinical Translation released by the International Society for Stem Cell Research (ISSCR). Human peri-gastruloid experiments followed a previously established protocol^65^ and were performed at UT Southwestern Medical Center. The UT Southwestern Stem Cell Oversight Committee (SCRO) conducted a comprehensive review and approval of the human peri-gastruloids protocol (registration no. 29) in accordance with the requisites stipulated by the 2021 ISSCR guidelines. Culture for these embryo models was limited to 8 days and was carefully calibrated to the minimal necessary time period for the examination of gastrulation and the incipient stages of organogenesis in human development. A significant point to emphasize is that these embryo models are designed to be incomplete due to their lack of trophoblast cells and are thus not capable of supporting full human development. No human embryos or gametes were used in this study.

### Cell lines and maintenance

Human primed embryonic stem cells (ESCs) (WIBR3 (OCT4-GFP) and UCLA4 (both female lines)^58, 164^), human induced pluripotent stem cells (hiPSCs) (CRISPRi (GEN1C clone, male)^57^), and chimp iPSCs (C6, C7)^59^ were cultured on Matrigel-coated dishes (Corning) in mTeSR1 medium (Stem Cell Technologies) at 37°C. Cells were passaged every 3-4 days using 0.02% EDTA solution (Sigma-Aldrich) for human ESCs and chimp iPSCs and Accutase (Life Technologies) for hiPSCs.

Naïve mouse ESCs (C57BL/6 x 129S4Sv/Jae F1-derived v6.5) were maintained at 37°C in naive mouse ESC medium composed of a 1:1 ratio of DMEM/F12 and Neurobasal medium supplemented with: 1X MEM non-essential amino acids, 1 mM sodium pyruvate, 2 mM L-glutamine, 100 U/mL penicillin, 100 μg/mL streptomycin, 50 μM β-mercaptoethanol, N2 and B27 supplements (all from Life Technologies), 1 μM PD0325901 (Axon Medchem), 3 μM CHIR99021 (Axon Medchem), and 10 ng/mL mouse leukemia inhibitory factor (mLIF) (R&D Systems).

### Cell line generation

GEN1C hiPSCs (2.5×10⁶ cells) were co-transfected with 7 μg of PB-mU6-sgRNA-CAG-mKate2-T2A-Blasticidin vector and 1.4 μg of transposase-encoding vector. The sgRNA sequences were: *ILF2* (CCTTAAAACACGAACAATGG) and *ILF3* (CGACAAGGTAAGCGTCAATG). Electroporation was performed using the Neon Transfection System (Life Technologies) at 1050 V, 30 ms, with 2 pulses. Stable transfectants were selected with Blasticidin beginning 4 days post-nucleofection for at least three passages, followed by sorting for mKate2-positive cells by FACS.

RUES2 hESCs (1×10⁶ cells) were co-transfected with 2 μg of AAVS1-Ndi-CRISPRi donor vector (modified from Addgene #73497 by replacing mCherry with EBFP2) and eSpCas9(1.1)_AAVS1_T2 vector (Addgene #79888). Electroporation was performed using previously established conditions. Stable integrants were identified by FACS selection of EBFP2-positive cells following doxycycline treatment. Selected clones were subsequently co-transfected with PB-RTTA-BsmBI-iRPF670 vector containing ILF3 sgRNA (CGACAAGGTAAGCGTCAATG) and PB-RTTA-BsmBI-GFP vector containing ILF2 sgRNA (CCTTAAAACACGAACAATGG), followed by FACS isolation of cells co-expressing both guide RNAs.

For the ILF3 mutant, rescue, and ADAR1 Co-IP experiments, HA-ILF3ΔRBM, HA-ILF3 WT, and FLAG-ADAR1 were individually cloned into doxycycline-inducible PiggyBac vectors with distinct antibiotic resistance markers (modified from PB_rtTA_BsmBI, Addgene #126028) and nucleofected into UCLA4 human ESCs. For rescue experiments, HA-ILF3 WT or HA-ILF3ΔRBM constructs were nucleofected into GEN1C CRISPRi sgILF3 cells. For co-immunoprecipitation studies, FLAG-ADAR1 was co-nucleofected with HA-ILF3 into wild-type GEN1C hiPSCs using the previously described electroporation parameters. For RNA immunoprecipitation followed by qPCR (RIP-qPCR), HA-ADAR1 was nucleofected into GEN1C wildtype cells.

For the ILF3 degron GEN1C cells, an FKBP12^F36V^-HA-2A-mCherry sequence was inserted into the first exon of ILF3 using CRISPR-Cas9-mediated homologous recombination. The donor plasmid was constructed by incorporating 500 bp ILF3-specific homology arms into pNQL004-SOX2-FKBPV-HA2-P2A-mCherry (Addgene #175552). An ILF3-targeting sgRNA (CACTACAGAAGAAGTAAAAA) was cloned into pSpCas9 (BB)-2A-Hygro (Addgene #127763). GEN1C hiPSCs (1×10⁶ cells) were co-nucleofected with 2µg of each plasmid using previous conditions. Single-cell clones expressing stable mCherry were isolated by FACS. Homozygous insertion was verified by PCR using primers spanning the ILF3 homology arms (Forward: AAAGATGTGTCCTGCTGTGT; Reverse: AGTAATGGGGCACTGACTTCG) and an mCherry-specific reverse primer (CCCCGTAATGCAGAAGAAGA). Western blot analysis confirmed ILF3 depletion upon dTAG^V^-1 treatment.

### Pluripotency exit assay

For the human pluripotency exit assay^1^, cells (4×10⁴) were seeded in Matrigel-coated 24-well plates in mTeSR1 medium (Stem Cell Technologies) supplemented with 10 μM Y-27632 (Axon Medchem). After 24 hours, the medium was replaced with differentiation conditions using either mTeSR1 Medium Without Select Factors (Stem Cell Technologies), maintained for 4 days, or mTeSR1 supplemented with 2.5 μM PD0325901 (Axon Medchem), maintained for 3 days followed by 24 hours in standard mTeSR1. OCT4-GFP or NANOG expression was analyzed by flow cytometry at the assay endpoint. For gene silencing experiments, genes were depleted either through shRNA infection of WIBR3 OCT4-GFP, UCLA4, or GEN1C wildtype cells 3 days pre-assay, or through doxycycline treatment (1 μg/mL, Sigma) of CRISPRi cells 3 days pre-assay.

For the chimp pluripotency assay, C6 and C7 chimp iPSCs were infected with shRNAs targeting *ILF2* or *ILF3* three days pre-assay (shRNAs used for human assays targeted sequences that were conserved in chimps). Cells were selected with puromycin, then seeded in Matrigel-coated 24-well plates in mTeSR1 medium (Stem Cell Technologies) supplemented with 10 μM Y-27632 (Axon Medchem). After 24 hours, the medium was replaced with mTeSR1 Medium Without Select Factors (Stem Cell Technologies) and maintained for either two or three days. NANOG expression was analyzed by flow cytometry at the assay endpoint.

For the mouse primed pluripotency exit assay, naïve mouse embryonic stem cells (2.74×10⁶) were seeded in Matrigel-coated (Corning) 6-well plates in N2B27 medium supplemented with 20 ng/mL Activin A (Peprotech), 12 ng/mL bFGF (Peprotech), and 1% KnockOut Serum Replacement (KSR, Life Technologies). After 2 days, primed ES cells were dissociated using Accutase (Life Technologies) for 5 minutes at room temperature, washed twice with DMEM-F12, and seeded (5×10⁴ cells per well) in Matrigel-coated (Corning) 24-well plates. After 24 hours, the medium was replaced with N2B27 supplemented with 20 ng/mL Activin A (Peprotech), 12 ng/mL bFGF (Peprotech), 1% KSR (Life Technologies), and 1 μM PD0325901 (Axon Medchem). NANOG expression was analyzed by flow cytometry after 48 hours. ILF2 and ILF3 were depleted either through shRNA infection of naïve cells in suspension 4 days pre-assay (day of primed induction), or through dTAG treatment (1 μM, or equivalent volume DMSO) of degron-tagged primed cells 3 days pre-assay (day after primed induction).

### ILF3 rescue experiments

GEN1C hiPSCs containing both the CRISPRi sgILF3 system and doxycycline-inducible constructs for either HA-ILF3 WT (GEN1C CRISPRi sgILF3/HA-ILF3 WT) or HA-ILF3ΔRBM (GEN1C CRISPRi sgILF3/HA-ILF3ΔRBM) were pretreated with 1 μg/mL doxycycline for 3 days to simultaneously downregulate endogenous ILF3 and express the exogenous ILF3 variants. Following pretreatment, cells underwent pluripotency exit, with NANOG expression evaluated by intracellular staining and flow cytometry after 4 days of culture in mTeSR1 supplemented with 2.5 μM PD0325901. GEN1C CRISPRi cells with a non-targeting guide and sgILF3 cells without rescue constructs served as controls. For RNA-seq and editing analysis, GEN1C CRISPRi sgILF3/HA-ILF3ΔRBM cells were pretreated with 1 μg/mL doxycycline for 3 days, then replated in mTeSR1 containing doxycycline for an additional 4 days before RNA sequencing.

To assess the downstream targets, rescue experiments were performed using GEN1C CRISPRi sgILF3 cells transduced with lentiviral vectors expressing BRD3, SMYD3, JARID2, or TEAD2 coding sequences under the control of the Ef1α promoter (Ef1α-IRES-Hygro), alongside a control vector. GEN1C sgCTRL cells were also infected with the control vector and served as a positive control for pluripotency exit. Following 3 days of hygromycin selection (75 μg/mL), cells were replated in mTeSR medium supplemented with 10 μM ROCK inhibitor (ROCKi) and 1 μg/mL doxycycline for 24 hours to induce ILF3 knockdown. ROCKi was then withdrawn, and doxycycline treatment was maintained for an additional 2 days. Subsequently, cells were seeded at a density of 40,000 cells per well in a 24-well plate in mTeSR medium. After 24 hours, the medium was replaced with mTeSR supplemented with 2.5 μM PD0325901 and 25 μg/mL hygromycin. RNA was collected after 3 days of PD treatment, and cells were fixed after 4 days for NANOG staining.

### Human two-dimensional gastruloids formation

GEN1C hiPSCs harboring CRISPRi sgCTRL or sgILF3 systems were treated with doxycycline (1 µg/mL) for 3 days to induce gene silencing. Cells were subsequently plated on laminin-521–coated (5 µg/mL) micropatterned plates (CYTOO) according to previously described protocols ^66^. Gastruloid formation was initiated by BMP4 treatment (50 ng/mL) for 48 hours.

### Human three-dimensional peri-gastruloids formation

Peri-gastruloids were generated from RUES2 hESCs harboring the CRISPRi sgILF2/sgILF3 system. Cells were first converted to expanded potential stem cells (hEPSCs) before differentiation into peri-gastruloids as previously described^65^. Briefly, EPSCs were dissociated into single cells with TrypLE, and feeder cells were removed through sequential 1-hour incubations on gelatin-coated plates. Single-cell EPSCs were then seeded into AggreWell800 plates (Stem Cell Technologies) at 150-180 cells per microwell in tHDM medium (N2B27 with 10 ng/mL FGF2 (Peprotech), 10 ng/mL Activin A (Peprotech), 1 µM CHIR99021 (Selleckchem), and 0.3 µM PD0325901 (Selleckchem)) supplemented with CEPT (50 nM Chroman 1 (Medchemexpress), 5 µM Emricasan (Selleckchem), 1 x Polyamine (Sigma-Aldrich), 0.7 µM *Trans*-ISRIB (Tocris)). On day 1, half of tHDM was replaced with tHDM without CEPT. On day 2, medium was changed to N2B27 supplemented with 10 ng/mL FGF2, 10 ng/mL Activin A, and 0.3 µM PD0325901 (tHDM without CHIR99021). On day 3, a half medium change was performed with tHDM without CHIR99021. On day 4, the medium was completely replaced with IVC1 medium (Advanced DMEM/F12, 1X Glutamax, 1X Insulin-Transferrin-Selenium-Ethanolamine (Gibco), 8 nM β-estradiol (Sigma-Aldrich), 200 ng/mL progesterone (Sigma-Aldrich), 25 µM N-acetyl-L-cysteine (Sigma-Aldrich), 1X sodium pyruvate, and 20% FBS) containing 4% Matrigel. On day 6, peri-gastruloids exhibiting embryonic disc-like structures and yolk sac cavities (∼60% of total) were selected and transferred to low-attachment plates in IVC2 medium (IVC1 medium, replacing 20% FBS with 30% knockout serum replacement (KSR, Thermo Scientific), and adjusting Advanced DMEM/F12 volume appropriately) with 4% Matrigel (15-20 structures per well). The structures were cultured under this condition until day 8.

### Mouse gastruloids formation

Gastruloids were generated from KH2 mouse embryonic stem cells following transduction with short hairpin RNAs (shRNAs) targeting either *Ilf2* or *Ilf3*, as previously described ^67^. Cells were seeded in ultra-low attachment round-bottom 96-well plates at a density of 300 cells per well in N2B27 medium. After 48 hours of culture, cells were treated with 3 µM CHIR99021 (Axon MedChem) for 24 hours. Morphological assessment of gastruloid elongation was performed at 5 days after seeding.

### Human neural progenitor cell culture and neuronal differentiation

iPSC-derived XCL-1 neural progenitor cells (NPCs, StemCell Technologies) were cultured on Matrigel-coated dishes at 37°C in STEMDiff Neural Progenitor Medium (StemCell Technologies).

For differentiation, NPCs were seeded at a density of 4.0 x 10^4^ cells/cm² and allowed to reach confluency over 4 days. Medium was then switched to DMEM/F12 (Sigma-Aldrich) supplemented with 1×N2 (Life Technologies), 1×B27 (Life Technologies), 300 ng/mL cAMP (Sigma-Aldrich), and 0.2 μM L-abscorbic acid (Sigma-Aldrich) for 6 days before analysis.

### Human myoblasts culture and differentiation

Primary human skeletal myoblasts (pooled male and female donors) were obtained from Thermo Fisher Scientific (Gibco, A12555) and cultured at 37°C under hypoxic conditions (5% O₂) in MegaCell DMEM (Sigma-Aldrich) supplemented with 5% fetal bovine serum (FBS, Lonza), 2 mM L-glutamine (Life Technologies), 0.1 mM β-mercaptoethanol (Life Technologies), 1× MEM Non-Essential Amino Acids Solution (Life Technologies), 5 ng/mL basic fibroblast growth factor (bFGF, PeproTech), 100 U/mL penicillin, and 100 μg/mL streptomycin.

For differentiation, myoblasts were seeded at a density of 1.6 × 10⁴ cells/cm², expanded, and allowed to spontaneously differentiate without passaging for 7 days, with medium changes every 2 days.

### Human definitive endoderm induction and maturation

GEN1C hiPSCs containing sgILF2, sgILF3, or a non-targeting control were sequentially differentiated into anterior primitive streak, definitive endoderm, and foregut as previously described^165^. Differentiation was performed using the following media compositions: on Day 1, cells were cultured in CDM2 base media supplemented with 100 ng/mL Activin A (Peprotech), 3 μM CHIR99021 (Axon MedChem), 20 ng/mL FGF2 (Peprotech), and 50 nM PI-103 (Tocris). On Day 2, CDM2 base media was supplemented with 100 ng/mL Activin A, 250 nM LDN-193189, and 50 nM PI-103. On Day 3, cells were cultured in CDM3 base media supplemented with 20 ng/mL FGF2, 30 ng/mL BMP4 (R&D Systems), 75 nM TTNPB (Selleck), and 1 μM A-83-01 (Reprocell). CDM2 basal medium consisted of 50% IMDM and 50% F12 with 2 mM L-glutamine (Sigma-Aldrich), penicillin-streptomycin (Sigma-Aldrich), 1 mg/mL polyvinyl alcohol (Sigma), 1% v/v chemically defined lipid concentrate (Thermo Fisher), 450 μM 1-thioglycerol (Sigma-Aldrich), 0.7 μg/mL recombinant human insulin (Sigma-Aldrich), and 15 μg/mL human transferrin (Sigma). CDM3 basal medium consisted of 45% IMDM and 45% F12 with 2 mM L-glutamine (Sigma-Aldrich), penicillin-streptomycin (Sigma-Aldrich), 10% KnockOut serum replacement (Thermo Fisher), 1 mg/mL polyvinyl alcohol (Sigma-Aldrich), and 1% v/v chemically defined lipid concentrate (Thermo Fisher).

### Virus production

HEK293T cells were co-transfected with transfer plasmid and packaging plasmids (VSV-G and D8.9 for lentiviral production) using calcium phosphate transfection. Viral supernatants were collected 24 hours post-transfection and concentrated by ultracentrifugation at 21,000 × g for 2 hours at 4°C. The concentrated virus was resuspended in Opti-MEM (Gibco) and stored at −80°C. Cells were transduced by adding the virus to the culture medium the day after passaging.

### Flow cytometry

For live cell analysis, DAPI or DRAQ7 was used as viability dye. For intracellular flow cytometry, cells were fixed with 4% paraformaldehyde and permeabilized with 0.2% Triton X-100 in PBS (0.1% PBST). Staining was carried out using NANOG (D73G4) XP Rabbit mAb (1:100, Alexa Fluor 647 conjugate, Cell Signaling 5448) or, for mouse-derived samples, mouse-specific NANOG (D2A3) XP Rabbit mAb (1:200, Cell Signaling 8822) followed by secondary antibody incubation with donkey anti-mouse Alexa Fluor 488 (1:500, Invitrogen A21202). Samples were acquired through FACSDiva software on an LSRII, LSRFortessa or FACSCanto II (BD Biosciences). Cells were analyzed using an LSRII, FACSCanto, or LSRFortessa flow cytometer (BD Biosciences) or sorted using a FACSAria (BD Biosciences). All flow cytometry data were analyzed using FACSDiva version 6.1.2 and FlowJo version 10 (BD Biosciences).

### Immunofluorescence

Cells were fixed with 4% paraformaldehyde, permeabilized with 100% methanol for ILF2 or ILF3 staining, or 0.2% Triton X-100 in PBS (0.1% PBST) for all other antibodies, blocked, and incubated overnight at 4°C with primary antibodies. The following day, cells were stained with Alexa Fluor-conjugated secondary antibodies, (1:500) Goat anti-Rabbit IgG (H+L) (Thermo Fisher) and Goat anti-Mouse IgG (H+L) (Thermo Fisher), for one hour at room temperature. Nuclear staining was performed using DAPI (BD Biosciences). The primary antibodies used in this study were ILF2 (1:200, Santa Cruz sc-365283), ILF3 (1:500, ProteinTech 19887-1-AP and 1:2000, A303-651A, Bethyl), NANOG (D73G4) XP Rabbit mAb (1:300, Cell Signaling 4903S), βIII-tubulin (TUJ1, 1:200, BioLegend 801211), SOX17 (1:100, R&D Systems AF1924), and MYOSIN (MF-20, Hybridoma Bank).

### Western blot

Samples were lysed in RIPA buffer (50 mM Tris-HCl, pH 8.0, 150 mM NaCl, 0.1% SDS, 0.5% sodium deoxycholate, 1% Triton X-100, 1 mM EDTA) supplemented with protease inhibitors and benzonase. Lysates were subjected to standard Western blotting procedures using the following primary antibodies: rabbit anti-human/mouse ILF2 (1:3000, Bethyl A303-147A), rabbit anti-human/mouse ILF3 (1:3000, Bethyl A303-651), anti-FLAG (1:2000, Addgene 194502), anti-HA (1:2000, BioLegend 901516), HRP-rabbit anti-human/mouse β-actin (1:3000, Cell Signaling 5125), and rabbit anti-human/mouse vinculin (1:2000, Cell Signaling 13901S). The following HRP-conjugated secondary antibodies were used: HRP-donkey anti-rabbit IgG (1:3000, BioLegend 406401), and HRP-horse anti-mouse IgG (1:3000, Cell Signaling 7076P2).

### Co-Immunoprecipitation (Co-IP)

For co-IP, frozen cell pellets (10x10^6^) were lysed in 500 μL of lysis buffer (50 mM Tris-HCl, pH 7.5, 150 mM NaCl, 1 mM EDTA, 1 mM EGTA, 5 mM MgCl₂, 0.5% Triton X-100) supplemented with fresh protease inhibitors (Roche) and incubated for 1 hour at 4°C with rotation. Lysates were pre-cleared by incubation with Protein G Mag Sepharose beads (Millipore Sigma) for 2 hours at 4°C.

For immunoprecipitation, fresh Protein G Mag Sepharose beads were blocked overnight in 1% BSA (Sigma-Aldrich) at 4°C, incubated with 6 µg ILF3 (Bethyl A303-651A) antibody or 6 µg rabbit IgG (Sigma-Aldrich, 12-370) for 2 hours at room temperature, and crosslinked using 20 mM DMP in 0.2 M triethanolamine for 30 minutes at room temperature. Pre-cleared lysates were then incubated with antibody-crosslinked beads overnight at 4°C. Beads were washed six times with cold lysis buffer containing protease inhibitors.

For SDS-PAGE and western blot validation, 5% of beads were boiled in Laemmli buffer (Bio-Rad) containing 5% 2-mercaptoethanol for 5 minutes. Western blot proceeded as previously described.

For FLAG-ADAR1 pulldown, immunoprecipitation was performed using 6 μg mouse anti-FLAG (Addgene 194502) or 6 μg rabbit IgG (Sigma-Aldrich 12-370).

### RNA Immunoprecipitation (RIP)

RNA immunoprecipitation was performed to assess protein-RNA interactions in native conditions. GEN1C wild-type hPSCs transfected with a doxycycline-inducible, HA-tagged ADAR1p110. They were then infected with either pLKO.1 shCTRL or pLKO.1 shILF3 and selected with 3µg/mL puromycin for two days. On the second day after infection, cells were treated with 2µg/mL doxycycline to induce HA-tagged ADAR1p110 expression. Three days after infection, cells were harvested and lysed in homogenization buffer (50 mM Tris pH 7.5, 100 mM KCl, 12 mM MgCl2, 1% IGEPAL, 1 mM DTT, 200 U/mL RNase inhibitor, 1 mg/mL heparin, 1% sodium deoxycholate, and protease inhibitors).

Fresh Protein G Mag Sepharose beads were blocked overnight in 1% BSA (Sigma-Aldrich) at 4°C, incubated with 2µg anti-HA antibody (BioLegend 901516) or IgG control for 2 hours at room temperature, and crosslinked using 20mM DMP in 0.2M triethanolamine for 30 minutes at room temperature. After centrifugation at 10,000 x g for 10 minutes at 4°C, clarified lysates were incubated with antibody-crosslinked beads overnight at 4°C with gentle rotation. Beads were washed four times with high-salt buffer (50 mM Tris pH 7.5, 300 mM KCl, 12 mM MgCl2, 1% IGEPAL, 0.5 mM DTT). RNA was directly isolated from the beads using RLT lysis buffer. For RNA-sequencing, libraries were prepared using the snapTotal-seq protocol as previously described^166^. For RT-qPCR analysis, RNA was reverse transcribed into cDNA using the LunaScript RT Mix (NEB). Enrichment was calculated as percent input and normalized to IgG control. All experiments were performed in biological triplicates.

### shRNA-mediated gene silencing

For constitutive gene silencing, oligonucleotide pairs encoding short hairpin RNAs (shRNAs) were annealed and cloned into either the pLKO.1-puro (Addgene #8453) or pSicoR-EF1α-mCherry-Puro (Addgene #31845) vectors. Knockdown efficiency was confirmed by RT-qPCR.

shRNA sequences used in this study:

- *ILF2* (pLKO backbone): CCTGGGATGGAGTGATAGTAA
- *ILF3* (pLKO backbone): CCAGAGGACGACAGTAAAGAA
- *ILF3* (pSicoR backbone): GGAGGTTGATGGCAATTCA
- *ADAR* (pSicoR backbone): GAACCCAAGTTCCAATACT
- *BRD3* (pSicoR backbone): GGGAGATGCTATCCAAGAA
- *SMYD3* (pSicoR backbone): GATCTGGAGTCAAATATTA
- *TEAD2* (pSicoR backbone): GAAATCCAGTCCAAGTTGA
- *JARID2* (pSicoR backbone): GCAAGTGACTGACCTCAAA
- Mouse *Ilf2* (pSicoR backbone): GAAACTGGCTTTGAAATTA
- Mouse *Ilf3* (pSicoR backbone): GGACTACACTGTTCAAATT

### UPF1 depletion experiment

GEN1C CRISPRi ILF3 degron cells were transduced with a Lentiguide mCherry Puro vector (Addgene #170510) targeting *UPF1* (AAAGTGAGAGTCTGCGAGCT). To induce *UPF1* silencing, cells were treated with 1 µg/mL doxycycline for 3 days. Following induction, cells were treated with DMSO, 1 µM dTAG^V^-1, or 1 µM dTAG^V^-1 combined with 1 µg/mL doxycycline for 24 hours. Cell pellets were then harvested for subsequent qPCR analysis.

### RNA preparation

Total RNA was isolated from cells for RT-qPCR using the Monarch Total RNA Miniprep Kit (NEB) and reverse transcribed into cDNA using the LunaScript RT Mix (NEB), following the manufacturer’s instructions. For RNA sequencing, RNA was extracted using the miRNeasy Mini Kit (Qiagen).

Chromatin-associated RNAs were isolated and extracted as previously described^121^. Briefly, 10⁷ cells were harvested, washed with cold PBS, and lysed in buffer containing 10 mM Tris pH 7.5, 150 mM NaCl, and 0.15% IGEPAL. Following centrifugation through sucrose buffer (10 mM Tris pH 7.5, 150 mM NaCl, 24% sucrose w/v) at 13,000 rpm for 10 min at 4°C, the cytoplasmic fraction was collected. Nuclear pellets were processed with glycerol buffer (20 mM Tris pH 7.9, 75 mM NaCl, 0.5 mM EDTA, 50% glycerol, 0.85 mM DTT) followed by nuclear lysis buffer (20 mM HEPES pH 7.6, 7.5 mM MgCl₂, 0.2 mM EDTA, 300 mM NaCl, 1 M urea, 1% IGEPAL, 1 mM DTT) to isolate the nucleoplasmic fraction (14,000 rpm, 2 min, 4°C). The remaining chromatin fraction was extracted using TRIzol and chloroform precipitation. All fractions were stored in RLT buffer before RNA purification using Qiagen RNeasy Mini columns with on-column DNase treatment.

### RT-qPCR analyses

Total RNA was extracted using the Monarch Total RNA Miniprep Kit (New England Biolabs) following the manufacturer’s instructions. Reverse transcription was performed using the LunaScript RT SuperMix Kit (New England Biolabs). For RT-qPCR, reactions were prepared with Luna Universal qPCR Master Mix (New England Biolabs) and pre-designed primers (Sigma-Aldrich) at a final concentration of 0.5 μM. qPCR was performed on a CFX96 Real-Time PCR Detection System (Bio-Rad) under the following cycling conditions: initial denaturation at 95°C for 1 minute, followed by 40 cycles of 95°C for 15 seconds and 60°C for 30 seconds. Primer sequences are available upon request.

### RNA-seq

Total RNA was extracted using the miRNeasy Micro Kit (QIAGEN) following the manufacturer’s instructions. For RNA-seq of ILF2, ILF3, and control CRISPRi, as well as ILF3 degron and NPCs experiments, polyadenylated RNAs were enriched, and cDNA libraries were generated using the NEBNext Ultra II Directional RNA Library Prep Kit for Illumina (New England Biolabs). For all other experiments, RNA-seq libraries were prepared using the snapTotal-seq protocol as previously described^166^. Libraries were sequenced on a NovaSeq X Plus (Illumina) with 150 bp paired-end reads.

### Assay for Transposase-Accessible Chromatin using sequencing (ATAC-seq)

ATAC-seq was performed as previously described^64^. Briefly, 50,000 cells were washed once with 100 μL PBS and resuspended in 50 μL lysis buffer (10 mM Tris-HCl, pH 7.4, 10 mM NaCl, 3 mM MgCl2, 0.2% IGEPAL). The nuclear suspension was centrifuged at 500 g for 10 minutes at 4°C, and the pellet was resuspended in 50 μL transposition reaction mix (25 μL TD buffer, 2.5 μL Tn5 transposase, and 22.5 μL nuclease-free water). The reaction was incubated at 37°C for 30 minutes. DNA was then isolated using the MiniElute Kit (QIAGEN), and libraries were amplified by PCR for 13 cycles. After amplification, libraries were size-selected for fragments between 100 and 1000 bp using AmpureXP beads (Beckman Coulter). The libraries were purified with the QIAquick PCR Purification Kit (QIAGEN) and assessed for integrity on a Bioanalyzer before sequencing.

### Cleavage Under Targets and Tagmentation (CUT&Tag)

CUT&Tag was performed as previously described^167^. Briefly, 60,000 bead-bound cells were permeabilized with Complete CT Antibody Buffer and incubated for 1h at room temperature on rotation with the following primary antibodies: anti-HA (1 µg, 13-2010, EpiCypher) anti-H3K27ac (0.5 µg, 39133, Active Motif) anti-H3K4me3 (0.5 µg, 39159, Active Motif), anti-H3K9me3 (0.5 µg, 39161, Active Motif), anti-H3K27me3 (0.5 µg, CST 9733S Cell Signaling Technology), and anti-H3K36me3 (0.5 µg, MA5-24687 Invitrogen). After incubation with secondary antibody (Guinea Pig anti-Rabbit IgG, ABIN101961), pAG-tethered transposase (pAG-Tn5 15-1017, Epicypher) was bound in situ at target loci. Following tagmentation and DNA clean-up, libraries were prepared by PCR amplification with barcoded Illumina-compatible primers. Libraries were assessed using the High Sensitivity D1000 ScreenTape (Agilent) before sequencing in paired-end 50 bp mode with Illumina NextSeq2000.

### Enhanced Crosslinking and Immunoprecipitation sequencing (eCLIP-seq)

2×10^7^ GEN1C cells per replicate were UV-crosslinked (400 mJ/cm²) to stabilize RBP-RNA interactions, then lysed. The whole cell lysates were sonicated and subjected to limited digestion with RNase I (40 U/mL of lysate), followed by immunoprecipitation for ILF2 or ILF3-RNA complexes using anti-ILF2 (10 µg Bethyl Laboratories A303-147A) or anti-ILF3 antibodies (10 µg Bethyl Laboratories A303-651A). Prior to immunoprecipitation, an aliquot of each extract was set aside and stored at 4°C to prepare the size-matched input control. Complexes were collected using anti-rabbit magnetic beads, washed, dephosphorylated, and 3’-ligated on-bead to custom oligonucleotides. All samples were run on 4-12% polyacrylamide gradient gels, and complexes were transferred to nitrocellulose membranes. Successful immunoprecipitation was confirmed by parallel Western blotting using the same anti-ILF2 or anti-ILF3 antibodies as described above. ILF2- or ILF3-RNA complexes were excised from the membrane and RNAs were released with proteinase K. Size-matched input samples were dephosphorylated and 3’-ligated. All samples were reverse-transcribed, and cDNAs were 5’-ligated on-bead, quantified by qPCR and PCR-amplified (< 18 cycles), followed by size selection (175-350 bp) on agarose gels and sequencing of libraries.

Remaining steps of eCLIP, including read processing and peak calling, were performed as previously described^168^ for two replicates of ILF2 and ILF3 as well as one paired size-matched input. Region-level analysis was performed as previously described^169^. The overlap between eCLIP peaks and Alu (RepeatMasker) were determined using Bedtools (version 2.30.0)^170^. Sense-oriented Alu elements located within gene bodies were annotated using RefSeq and UCSC Known Genes. Enrichment of eCLIP signal over these sense Alu elements was quantified using deepTools (v3.5.4^171^).

### Co-IP and Mass Spectrometry

Co-immunoprecipitation (Co-IP) of ILF3 and ILF2 was performed as previously described (see above) using 6 µg rabbit anti-ILF3 antibody (Bethyl Laboratories, A303-651A), 6 µg rabbit anti-NF45 antibody (Bethyl Laboratories, A303-147A), or 6 µg rabbit IgG (Sigma-Aldrich, 12-370). Following immunoprecipitation, the bulk Co-IP beads were analyzed by mass spectrometry. The proteins were reduced, alkylated with iodoacetamide, and digested separately with trypsin overnight. The resultant peptides were quantified using the Quantitative Fluorometric Peptide Assay (#23290, Thermo Fisher Scientific), and sample quantities were normalized.

For each sample, 125 ng of peptides were loaded onto a column and analyzed using a 90-minute gradient LC-MS/MS method on an Orbitrap Fusion Tribrid Mass Spectrometer (Thermo Fisher Scientific). The resulting LC-MS data were processed in Proteome Discoverer 2.5, and proteins were identified utilizing the SEQUEST search algorithm by searching against a human database and a database of common contaminants. The Minora node was used to perform Label-Free Quantification (LFQ) using the MS peak areas for each of the peptide-spectral matches (PSMs). The false discovery rate (FDR) for proteins and peptides was calculated using the Protein FDR validator and Percolator nodes, respectively, employing a target-decoy approach. Peptides with an FDR of <1% were included, and all proteins were included if identified with at least two peptides. Data were analyzed using R.

### LC-MS/MS Proteomics

hiPSCs GEN1C cells were collected after 0, 1, and 4 days of 1 mM dTAG^V^-1 treatment. Cells were resuspended in 100 µL of 5.4 M guanidine hydrochloride in 100 mM Tris-HCl, pH 8.0, and gently vortexed at ambient temperature for at least 20 minutes. A bicinchoninic acid (BCA) protein assay (Pierce) was performed according to the manufacturer’s instructions, after which sufficient guanidine hydrochloride in 100 mM Tris-HCl, pH 8.0 was added to bring the protein concentration to 1 mg/mL. Next, 50 µg of protein from each cell pellet was transferred into separate 1.5-mL microcentrifuge tubes. Samples were incubated in a sand bath at 110 °C for 5 minutes, then cooled at room temperature for 5 minutes, followed by a second incubation at 110 °C for 5 minutes. Then, 450 µL of LC-MS-grade methanol was added to each sample and vortexed for 10 seconds. The samples were centrifuged at 14,000 x g for 2 minutes at 4 °C to pellet the protein. After carefully discarding the supernatant, each pellet was resuspended in 50 µL of lysis buffer (8 M urea, 100 mM Tris-HCl, pH 8.0, 10 mM TCEP, and 40 mM 2-chloroacetamide) and vortexed for 10 minutes at room temperature to resolubilize the protein. Then, 1 µL of 1 mg/mL LysC (VWR) prepared according to the manufacturer’s instructions was added to each sample, and the samples were incubated at room temperature for 4 hours with gentle rocking. Afterward, the samples were diluted with freshly prepared 100 mM Tris-HCl, pH 8.0, to reach a final urea concentration of 2 M, and 2.5 µL of 0.4 mg/mL trypsin (Promega) was added. Samples were incubated overnight at room temperature with gentle rocking. To stop the digestion, 40 µL of 10% TFA in water was added to each sample, followed by centrifugation at 14,000g for 2 minutes to pellet any insoluble material. The supernatants were desalted using Sep-Pak C18 cartridges (Waters). The desalted peptides were dried in a vacuum centrifuge (Thermo Fisher Scientific). The dried peptides were resuspended in 0.2% formic acid in water, and peptide concentrations were determined with a NanoDrop One Microvolume UV-Vis spectrophotometer (Thermo Fisher Scientific). The samples were then vialed and placed into a 5 °C autosampler for mass spectrometry analysis.

### RNA-seq analysis

Raw sequencing files (FASTQ format) were aligned to the human reference genome (GRCh37) or mouse reference genome (GRCm39) using STAR (version 2.5.1b^172^) with default settings. Duplicate reads were removed using samtools (version 1.3.1^173^), and gene-level read counts were generated using featureCounts (version 2.0.2^174^). Differential expression analysis was conducted with the R package DESeq2^175^. Genes were considered differentially expressed if they met the criteria of log2 fold change |(log2FC)| > 0.58 and adjusted P value < 0.05. Enrichment analysis of functional categories was performed using EnrichR^176^. Normalized bigWig files for genome browser visualization were created with Deeptools (version 3.5.4^171^).

A-to-I RNA editing events were identified using JACUSA2 (version 2.0.2^177^), retaining sites with read coverage ≥ 10. The enrichment of editing events in eCLIP-seq peaks was calculated with regioneR^178^. Editing ratios were calculated as the number of A-to-I editing events divided by total read coverage at the editing site and visualized using ggplot2 (version 3.4.0).

Differential alternative splicing was quantified using rMATS (version 4.1.1^179^) with the non-default parameters --readLength 151 --novelSS. Splicing events were considered significant if they met the criteria of FDR < 0.05 and |ΔPSI| ≥ 0.1. JET analysis was performed following a previously described pipeline^122^. The enrichment of JETs in transposable elements (RepeatMasker) was calculated using the R package regioneR^178^. For analysis of Alu orientation around included exons, RepeatMasker annotations were obtained from the UCSC Genome Browser. BEDtools^170^ was used to identify sequences within 500 base pairs flanking differentially spliced exons. Complementary Alu pairs were defined as at least one plus-stranded and one minus-stranded Alu family element located on opposite sides of the exon. The frequency of exons containing complementary versus non-complementary Alu pairs was quantified. Statistical significance was assessed by comparing these frequencies to those of differentially spliced exons without paired Alu elements using χ² test.

Premature termination codon analysis and differential isoform fraction calculation with DEXSeq method was performed using IsoformSwitchAnalyzeR (version 1.16.0)^180^. Premature termination codons (PTCs) were identified as stop codons occurring ≥50 nucleotides upstream of the canonical termination site in the reference annotation.

### ATAC-Seq analysis

Raw sequencing reads were trimmed using cutadapt (version 1.16) to remove adaptor sequences and then mapped to the human reference genome (GRCh37) using Bowtie2 (version 2.2.8). Duplicate reads were removed using Picard (http://broadinstitute.github.io/picard/). Reads that were not mapped to chromosomes or that had low-quality scores were discarded using samtools (version 1.3.1^181^). Peak calling was performed using MACS2 (version 2.2.9.1^182^). Differentially accessible peaks were identified using the R package csaw (version 1.28.0^183^) defined as peaks with p < 0.05 and log2FC > 0.58. Normalized bigwig files for visualization were generated using Deeptools (version 3.5.4^171^).

### CUT&Tag analysis

Sequencing reads were mapped to hg38 reference genome using Bowtie2^184^. Peak calling was performed using SEACR^79^, selecting the top 1% of regions based on AUC (stringent; 0.01). Heatmaps and average profiles of CUT&Tag read densities were generated using deepTools^171^. Bedtools was used to calculate read densities over identified peaks^170^. Based on RPKM distribution, a filter for RPKM (>0.5) was applied for HA peaks. Gene Ontology analysis was performed with ClusterProfiler after annotation using ChipSeeker^185, 186^. Plots were generated in R using the ‘ggplot2’ package.

### RIP-seq analysis

Mouse ILF3 RIP-seq data^102^ was downloaded from GEO (accession: GSE95145). Raw sequencing reads were mapped to the mouse reference genome (GRCm39) using Bowtie2 (version 2.2.8). Duplicates were removed using Picard (http://broadinstitute.github.io/picard/). Reads with low-quality were filtered using samtools (version 1.3.1^181^). Peak calling was performed using MACS2 (version 2.2.9.1)^182^ with the --no_lambda and --no_model options. Peaks present across replicates were obtained using Bedtools (version 2.30.0)^170^. The enrichment of transposable elements (RepeatMasker mm39) in RIP-seq peaks was calculated using a permutation test with the R package regioneR.

For human ILF3 RIP-seq samples, raw sequencing files were aligned to the human reference genome (GRCh37) using Bowtie2 (version 2.2.8) with the following parameters: --very-sensitive --local --no-discordant --no-mixed. Peak calling was performed using MACS2 (version 2.2.9.1) ^182^, with peaks called separately for each replicate, treating IP samples as experimental data and input RNA as control data. Peak lists were filtered for those that were present in both replicates, resulting in condition-specific peak sets (ADAR1 IP + shCTRL, ADAR1 IP + shILF3, IgG IP + shCTRL). IgG IP peaks were then subtracted from ADAR1 IP peaks using bedtools (version 2.30.0)^170^ – peaks with a greater than 90% overlap with IgG peaks were completely removed, while partial overlaps were trimmed. The final peak analysis included only peaks ≥40 nucleotides in length and located within reference chromosomes (chr1-chr22, X, Y; excluding chrM) and transcripts (Gencode). Peak annotation was performed using ChIPseeker (version 1.30.3) with GRCh37 annotations, prioritizing genomic features in the following order: 5’UTR, 3’UTR, exon, intron, downstream, and intergenic regions. Quantification of RIP-seq reads were performed with StringTie (version 2.2.3)^187^, followed by normalization using DESeq2.

### Proteomics analysis

Sample analysis was performed using a Vanquish Neo UHPLC system (Thermo Fisher Scientific) coupled to an Orbitrap Ascend Tribrid mass spectrometer (Thermo Fisher Scientific). Mobile phase A was water with 0.2% formic acid, and mobile phase B was 80:20 v/v acetonitrile:water with 0.2% formic acid. The gradient elution was carried out at a flow rate of 0.300 µl/min. Peptides (750 ng) were loaded onto a 75-µm i.d. column packed with 1.7-µm, 130 Å pore size BEH C18 particles (Waters Corp.), and the column was heated to 50°C during analysis. MS1 scans were acquired from 0 to 80 minutes at a scan range of 300–1,350 m/z, with a resolution of 240,000, a maximum injection time of 50 ms, and a normalized AGC target of 250%, corresponding to 1 × 10^6^ ions. Precursor ions were isolated from a 0.8-Da window, and data-dependent HCD MS2 scans were performed with 23% normalized collision energy and a normalized AGC target of 250%, equivalent to 2.5 × 10⁴ ions. MS2 scans were collected in the ion trap from 150–1,350 m/z with a maximum injection time of 12 ms and a dynamic exclusion period of 20 s.

Raw proteomic data files were processed using MSFragger v.4.1^188, 189^. The UniProt reviewed human protein database was retrieved on Jan 17, 2024 and used for protein identification. Label-free quantification was performed with a minimum ratio of 1, and the “match between runs” option was enabled. Protein groups with an intensity value of zero in 50% or more of the analyzed samples were removed. Missing quantitative values in the remaining proteins were imputed, log2-transformed, and subjected to statistical analysis using Argonaut3^190^.

### Alphafold3 modeling of human ILF2/3 complex-bound Alu dsRNA

The core regions of human ILF2 (amino acids 1-390) and ILF3 (amino acids 1-702) were modeled in complex with a 70-nucleotide Alu RNA sequence (5’-CGAGACGGGGUUUCACCAUGUUGGCCAGGCUGGUCUCGAACUCCUGACCUC- AGGUGAUCCGCCCGCCUCG-3’). The top five structural predictions were generated and compared, demonstrating a high degree of similarity across models. The highest-confidence prediction was selected for visualization using PyMOL (version 3.1). Although the complete protein sequences were included in the modeling predictions, visualization was focused on structured domains for clarity: the DZF domain (amino acids 5-378), dsRBM1 (amino acids 398-467), and dsRBM2 (amino acids 524-590) for ILF3, and the DZF domain (amino acids 24-370) for ILF2. Unstructured regions were hidden in the final representation.

### Statistical analysis

Quantitative data are expressed as mean ± standard deviation (s.d.). Statistical analyses were performed using Prism 10.2.2 software (GraphPad). Specific details of the statistical analyses, including the number of replicates, are provided in the figure legends. Bioinformatics analyses and statistical computations were conducted using R 4.1.0. Boxplots represent the mean along with the quartiles, with whiskers indicating the minimum and maximum quartiles, excluding outliers.

---

**Extended Data Figure 1.**
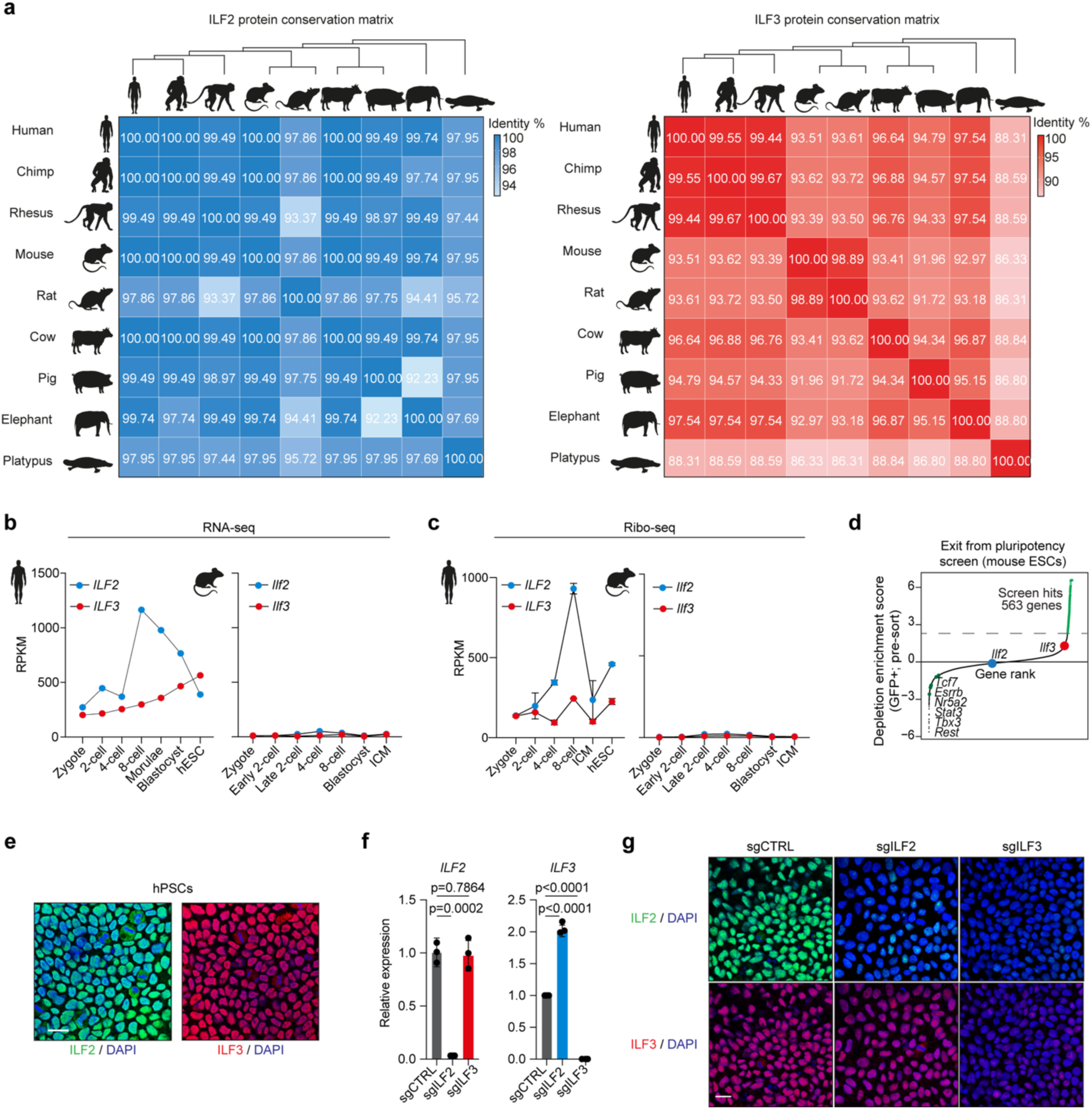
ILF2 and ILF3 protein sequences are conserved across species. (**a**) Matrices showing conservation of ILF2 (left) and ILF3 (right) proteins across species. (**b**) *ILF2* and *ILF3* expression from RNA-seq data in human^53^ and mouse^54^ embryos. (**c**) *ILF2* and *ILF3* expression from Ribo-seq data in human^55^ and mouse^54^ embryos. (**d**) Results from a mouse ESC exit screen after two days of differentiation, depicting depletion-enrichment scores^43^. (**e**) Representative immunofluorescence images of ILF2 (green) and ILF3 (red) localization in GEN1C hPSCs with DAPI nuclear counterstain (blue). Scale bar, 20 µm. (**f**) RT-qPCR quantification of *ILF2* and *ILF3* expression in control and CRISPRi-targeted cells (n = 3 biological replicates; mean ± s.d.; P values determined by one-way ANOVA with Tukey’s multiple comparisons test). (**g**) Representative immunofluorescence images of ILF2 (green) and ILF3 (red) localization in GEN1C CRISPRi hPSCs after 3 days of doxycycline treatment with DAPI nuclear counterstain (blue). Scale bar, 20 µm.

**Extended Data Figure 2.**
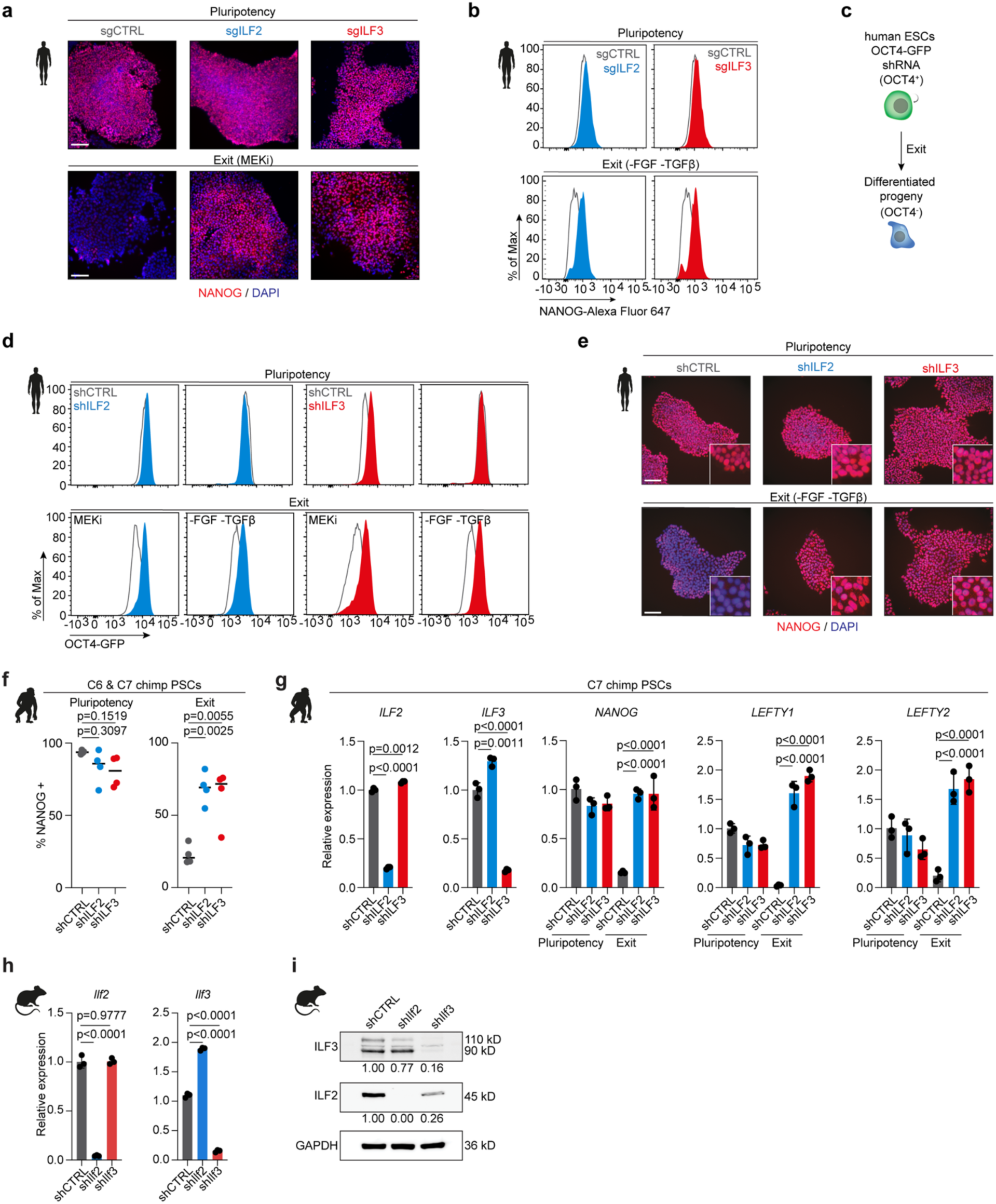
ILF2 and ILF3 are required for primate, but not mouse pluripotency exit. (**a**) Immunofluorescence analysis of NANOG expression in control and *ILF2* or *ILF3*-depleted hPSCs under self-renewal and exit conditions by MAPK pathway inhibition. Scale bar, 100µm. (**b**) Representative flow cytometry quantification of NANOG in hPSCs under self-renewal and exit conditions bFGF/TGFβ-depletion following *ILF2* or *ILF3* knockdown. (**c**) Schematic of differentiation protocol for WIBR3-OCT4GFP human ESCs. (**d**) Flow cytometry analysis of OCT4-GFP expression in cells cultured in mTeSR1, MAPK pathway inhibition, or bFGF/TGFβ-depleted conditions. (**e**) Immunofluorescence analysis of NANOG expression (red) with DAPI nuclear counterstain (blue) in cells cultured in mTeSR1 or bFGF/TGFβ-depleted conditions. Scale bar, 100 µm. (**f**) Quantification of NANOG-positive cells under self-renewal and exit conditions following *ILF2* or *ILF3* depletion in chimpanzee PSCs (n = 4 biological replicates; P values determined by one-way ANOVA with Tukey’s multiple comparisons test). (**g**) RT-qPCR analysis of chimpanzee PSCs under self-renewal or differentiation conditions following *ILF2* or *ILF3* depletion (n = 3 biological replicates; mean ± s.d.; P values determined by one-way ANOVA with Tukey’s multiple comparisons test). (**h**) RT-qPCR analysis of *Ilf2* and *Ilf3* expression in mouse embryonic stem cells (n = 3 biological replicates; mean ± s.d.; P values determined by one-way ANOVA with Tukey’s multiple comparisons test). (**i**) Representative western blot with signal quantification depicting validation of ILF2 and ILF3 knockdown efficiency in mESCs expressing shRNAs.

**Extended Data Figure 3.**
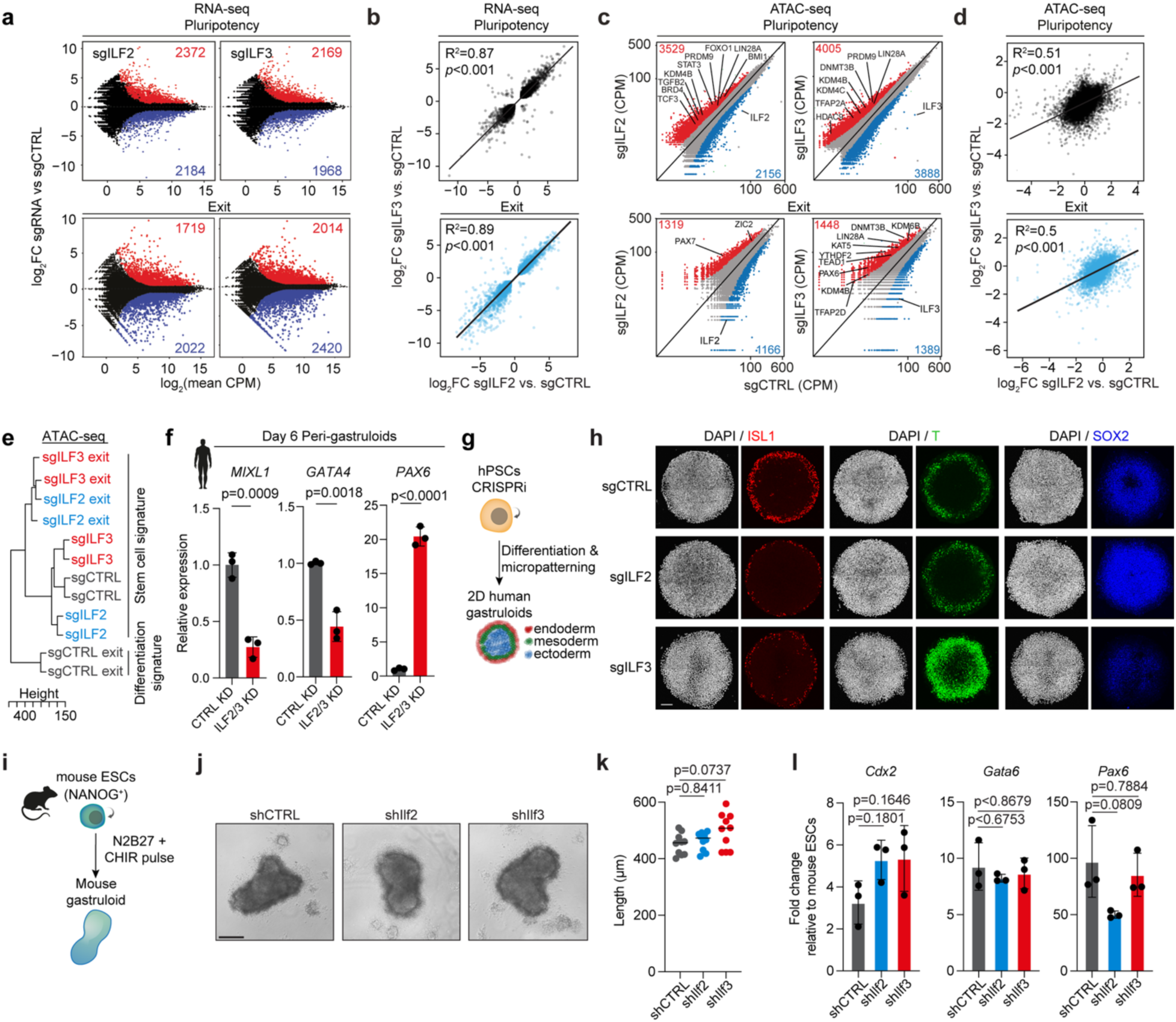
*ILF2* and *ILF3* suppression disrupts the transcriptional and chromatin profiles of hPSCs. (**a**) Differential gene expression analysis in *ILF2*- or *ILF3*-depleted cells under self-renewal and exit conditions (n = 2 biological replicates per condition; |FC| > 1.5; P < 0.05; red: upregulated, blue: downregulated). (**b**) Correlation analysis of transcriptional changes between *ILF2*- and *ILF3*-depleted cells under self-renewal and exit conditions. (**c**) ATAC-seq analysis of chromatin accessibility changes in *ILF2*- or *ILF3*-depleted cells (n = 2 biological replicates; |FC| > 1.5; P < 0.05; red: increased accessibility, blue: decreased accessibility). (**d**) Correlation analysis of chromatin accessibility changes between *ILF2*- and *ILF3*-depleted cells under self-renewal and exit conditions. (**e**) Hierarchical clustering of ATAC-seq datasets from hPSCs in self-renewal and exit conditions. (**f**) RT-qPCR analysis of differentiation markers in control and *ILF2/3*-depleted human peri-gastruloids (n = 3 biological replicates; mean ± s.d.; P values determined by one-way ANOVA with Tukey’s multiple comparisons test). (**g**) Experimental schematic for two-dimensional human gastruloid formation. (**h**) Representative immunofluorescence analysis of germ layer markers in control and *ILF2* or *ILF3*-depleted 2D human gastruloids. Scale bar, 100 µm. (**i**) Schematic of mouse gastruloid formation experimental design. (**j**) Representative brightfield images of *Ilf2*- and *Ilf3*-depleted mouse gastruloids. Scale bar, 200 µm. (**k**) Length analysis of *Ilf2*- and *Ilf3*-depleted mouse gastruloids (n = 10 biological replicates; mean ± s.d; P values determined by one-way ANOVA with Tukey’s multiple comparisons test). (**l**) RT-qPCR analysis of *Ilf2*- and *Ilf3*-depleted mouse ESCs (n = 3 biological replicates; mean ± s.d.; P values determined by one-way ANOVA with Tukey’s multiple comparisons test).

**Extended Data Figure 4.**
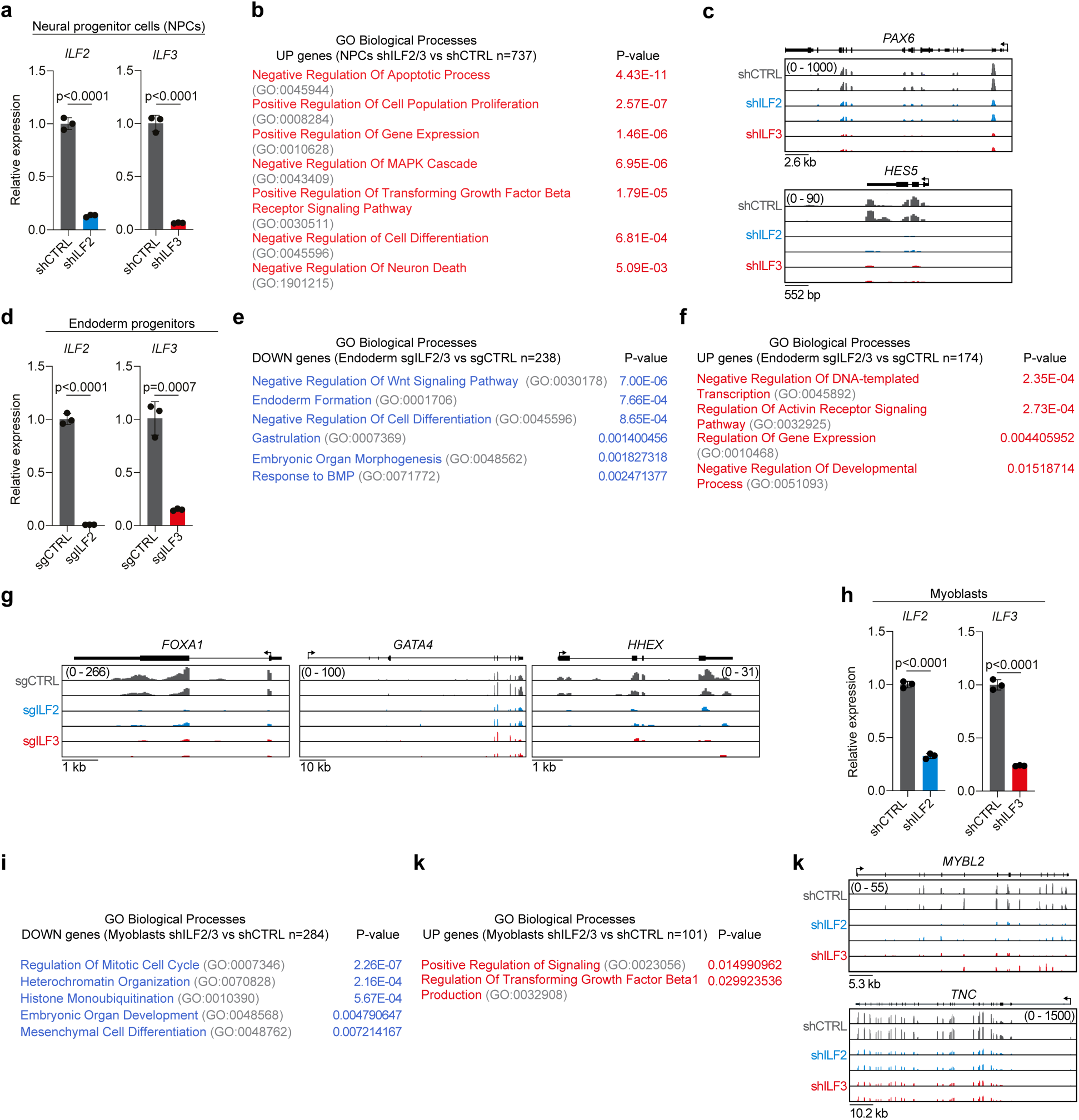
ILF2 and ILF3 regulate lineage-specific gene expression programs during progenitor cell differentiation. (**a**) RT-qPCR validation of *ILF2* and *ILF3* knockdown efficiency in neural progenitor cells (NPCs) (n = 3 biological replicates; mean ± s.d.; P values determined by unpaired Student’s t-test). (**b**) Gene Ontology analysis of upregulated genes in *ILF2*- or *ILF3*-depleted NPCs during differentiation (fold change > 1.5; P < 0.05; two-tailed Fisher’s exact test). (**c**) Representative RNA-seq tracks showing expression changes of individual genes in NPCs. (**d**) RT-qPCR validation of CRISPRi-mediated *ILF2* and *ILF3* silencing in endoderm progenitors (n = 3 biological replicates; mean ± s.d.; P values determined by unpaired Student’s t-test). (**e**) Gene Ontology analysis of downregulated genes in *ILF2*-or *ILF3*-depleted endoderm progenitors (fold change < -1.5; P < 0.05; two-tailed Fisher’s exact test). (**f**) Gene Ontology analysis of upregulated genes in *ILF2*- or *ILF3*-depleted endoderm progenitors (fold change > 1.5; P < 0.05; two-tailed Fisher’s exact test). (**g**) Representative RNA-seq tracks showing expression changes of individual genes in endoderm progenitors. (**h**) RT-qPCR validation of *ILF2* and *ILF3* knockdown efficiency in primary human myoblasts (n = 3 biological replicates; mean ± s.d.; P values determined by unpaired Student’s t-test). (**i**) Gene Ontology analysis of downregulated genes in *ILF2*-or *ILF3*-depleted myoblasts (fold change < -1.5; P < 0.05; two-tailed Fisher’s exact test). (**j**) Gene Ontology analysis of upregulated genes in *ILF2*- or *ILF3*-depleted myoblasts (fold change > 1.5; P < 0.05; two-tailed Fisher’s exact test). (**k**) Representative RNA-seq tracks showing expression changes of individual genes in myoblasts.

**Extended Data Figure 5.**
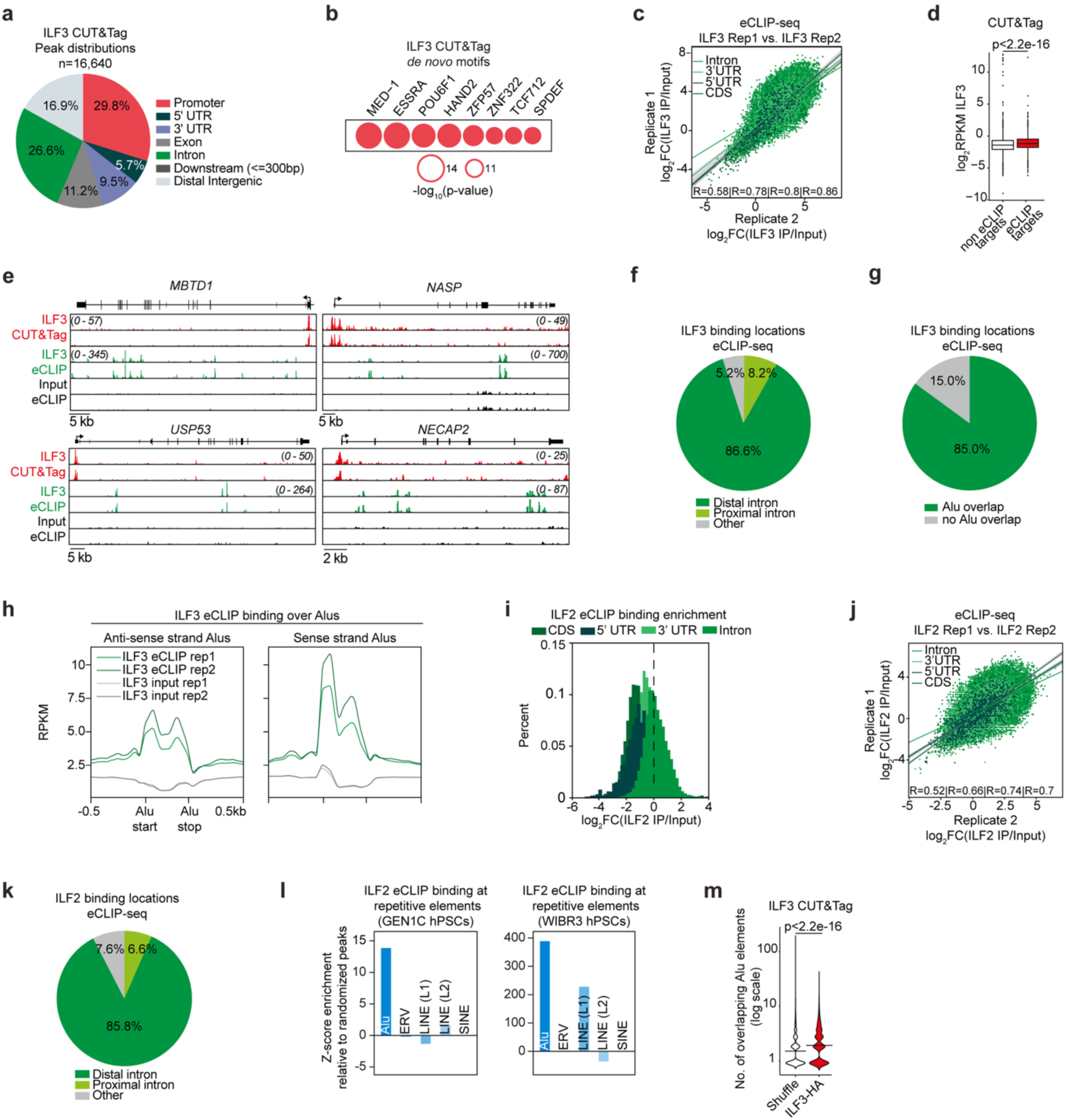
ILF2 and ILF3 bind chromatin and RNA at Alu elements. (**a**) Distribution of ILF3 CUT&Tag binding sites across genomic regions (RPKM>0.5; top 1% of peaks by area under the curve (AUC)). (**b**) De novo motif analysis of ILF3 CUT&Tag binding sites using HOMER. (**c**) Correlation analysis of ILF3 eCLIP signal indicating region-based log_2_FC enrichment between biological replicates (Correlation coefficients shown for intron, 3’UTR, 5’UTR, and CDS). (**d**) ILF3 chromatin binding intensity at regions overlapping and non-overlapping with eCLIP targets (P values determined by Wilcoxon rank-sum test). (**e**) Representative CUT&Tag and eCLIP tracks showing ILF3 targets. (**f**) Distribution of ILF3 eCLIP binding sites across genomic regions (log₂FC > 2.5, P < 0.05). (**g**) Distribution of ILF3 eCLIP binding sites that overlap with Alu elements (log₂FC > 2.5, P < 0.05). (**h**) Distribution of ILF3 eCLIP signal centered around sense and anti-sense Alu elements. (**i**) Distribution of ILF2 eCLIP signal enrichment relative to size-matched input controls (log₂FC > 3, P < 0.05). (**j**) Correlation analysis of ILF2 eCLIP signal log₂FC enrichment between biological replicates across genomic features. (**k**) Distribution of ILF2 eCLIP binding sites across genomic regions (log₂FC > 2.5, P < 0.05). (**l**) Enrichment analysis of transposable element families in ILF2 eCLIP targets from GEN1C hPSCs (left) and WIBR3 human ESCs (right) relative to randomized peak distributions. (**m**) ILF3 CUT&Tag binding versus shuffled peaks at Alu elements (P values determined by Wilcoxon rank-sum test).

**Extended Data Figure 6.**
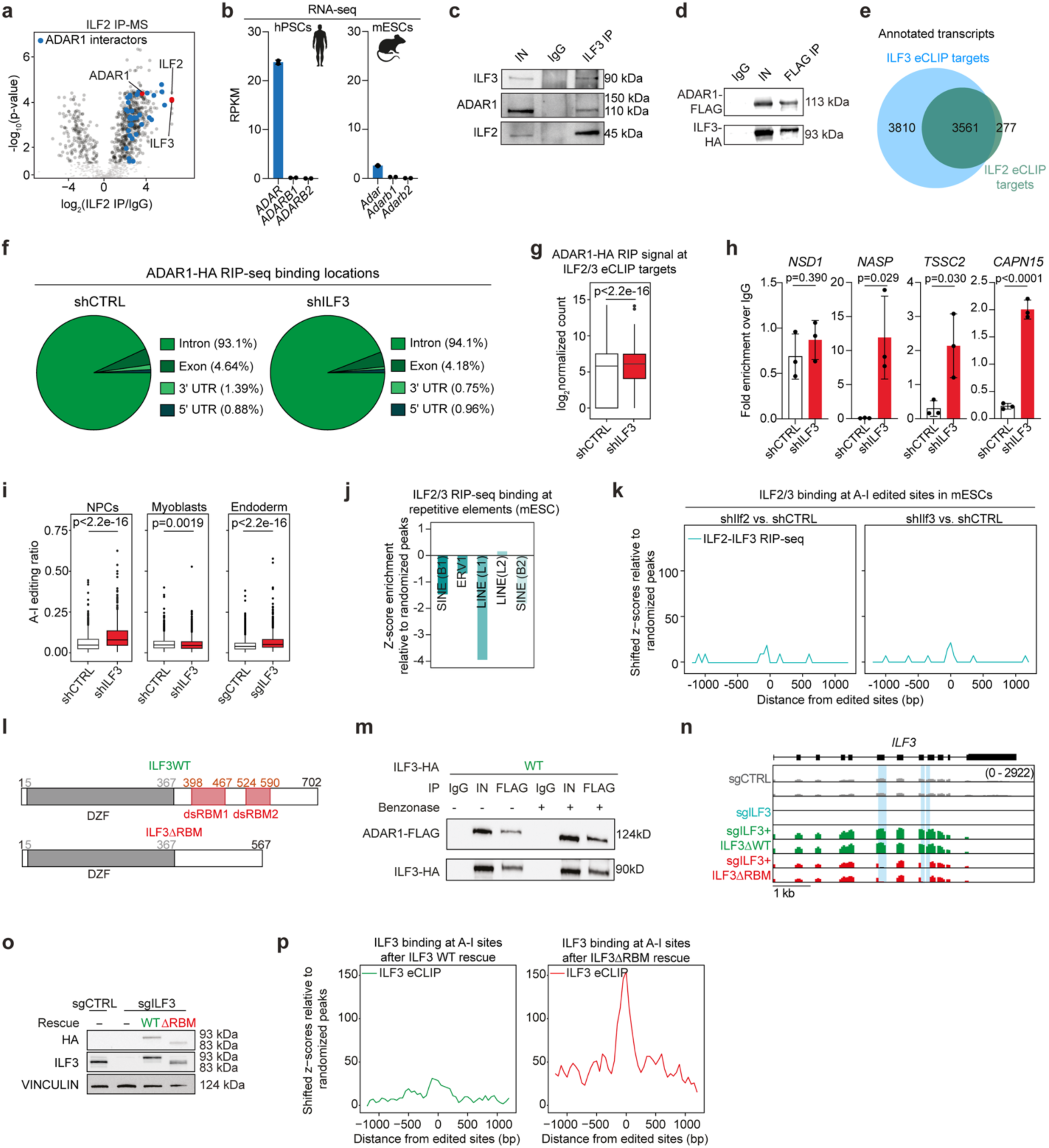
ILF2 and ILF3 inhibit RNA editing by ADAR1 in human adult progenitor cells. (**a**) Proteomic analysis of ILF2 interactors (n = 3 biological replicates). (**b**) Abundance of ADAR family transcripts in hPSCs (left) and mESCs (right). (**c**) Co-immunoprecipitation analysis showing ILF3-ADAR1 interaction after ILF3 pull-down. (**d**) Co-immunoprecipitation analysis showing ILF3-ADAR1 interaction after FLAG-ADAR1 pull- down in an alternative hPSC line (UCLA4). (**e**) Overlap between ILF2 and ILF3 eCLIP-bound transcripts. (**f**) Distribution of ADAR1-HA RIP-seq binding sites across genomic regions after control or *ILF3* knockdown (P < 0.0001). (**g**) ADAR1-HA binding strength to ILF2/3 eCLIP targets after control or *ILF3* knockdown (P value determined by Wilcoxon rank-sum test). (**h**) RT-qPCR of ADAR1-bound RNAs, normalized to input, before and after *ILF3* depletion (n = 3 biological replicates; mean ± s.d.; P values determined by unpaired Student’s t-test). (**i**) A-to-I editing frequencies in adult progenitor cells following *ILF3* knockdown (P values determined by Wilcoxon rank-sum test). (**j**) Transposable element family enrichment in ILF2/3 RIP-seq targets from mouse ESCs^102^ relative to a randomized peak distribution. (**k**) Aggregate plot showing ILF2 and ILF3 RIP-seq signal distribution^102^ centered around A-to-I-edited sites after *Ilf2* (left) and *Ilf3* (right) knockdown in mouse ESCs. Z-scores calculated relative to a randomized peak distribution. (**l**) Domain organization of wild-type and ΔRBM mutant ILF3. (**m**) Western blot analysis of FLAG co-immunoprecipitation with and without benzonase treatment. (**n**) Representative RNA-seq tracks showing expression changes of *ILF3* in CRISPRi sgCTRL, sgILF3, and rescue cells. Highlighted regions encode the double-stranded RNA-binding motifs. (**o**) Western blot validation of *ILF3* knockdown and rescue with wild-type or ΔRBM mutant ILF3. (**p**) Aggregate plot showing ILF3 eCLIP signal distribution centered around A-to-I-edited sites after *ILF3* knockdown followed by rescue with ILF3 WT (left) or ΔRBM mutant (right). Z-scores calculated relative to a randomized peak distribution.

**Extended Data Figure 7.**
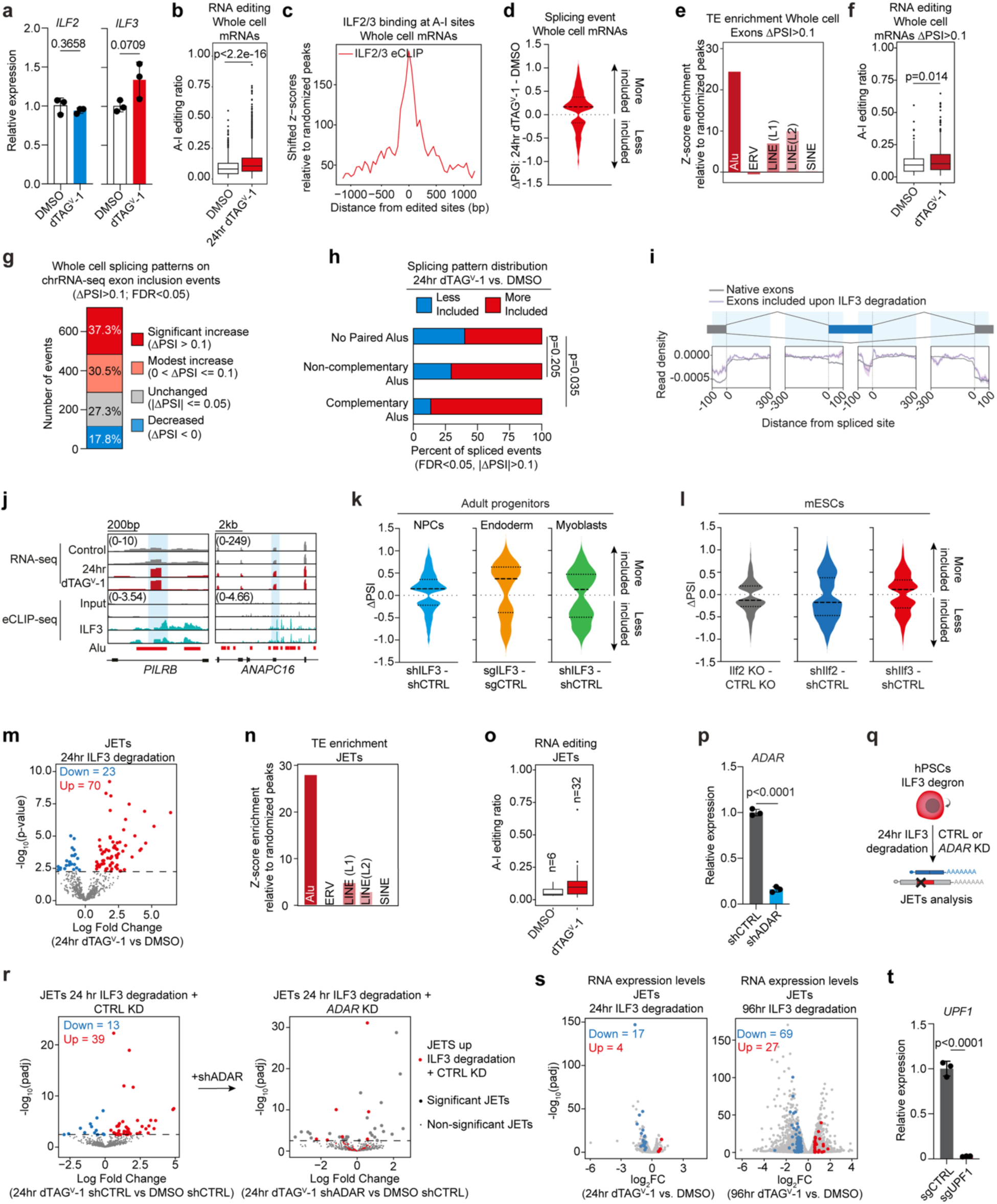
ILF3 degradation causes widespread mis-splicing. (**a**) RT-qPCR analysis of *ILF2* and *ILF3* expression in ILF3 degron hPSCs treated with DMSO or dTAG^V^-1 for 24 hours (n = 3 biological replicates; mean ± s.d.; P values determined by unpaired Student’s t-test). (**b**) Editing frequencies in whole cell-extracted mRNAs after 24 hours of ILF3 degradation (n = 2 biological replicates; P values determined by Wilcoxon rank-sum test). (**c**) Aggregate plot showing ILF3 eCLIP signal distribution centered around A-to-I-edited sites in whole cell-extracted mRNAs after 24 hours of ILF3 degradation. Z-scores calculated relative to a randomized peak distribution. (**d**) Alternative splicing events in whole cell-extracted mRNAs 24 hours after ILF3 degradation (|ΔPSI| > 0.1, FDR < 0.05). (**e**) Enrichment of repetitive elements in cassette exons more included after ILF3 degradation in whole cell-extracted mRNAs (ΔPSI > 0.1, FDR < 0.05) relative to a randomized peak distribution. (**f**) Editing frequencies in transcripts showing increased exon inclusion after ILF3 degradation (ΔPSI > 0.1, FDR < 0.05; n = 2 biological replicates; P values determined by Wilcoxon rank-sum test). (**g**) Distribution of whole cell RNA splicing patterns at chromatin-associated exonization events after 24 hours of ILF3 degradation. Categories based on events with increased exon inclusion in chromatin-associated mRNAs (ΔPSI > 0.1, FDR < 0.05). (**h**) Categorization of significant cassette exon splicing events based on Alu element configuration within ±500 bp flanking regions. ‘No Paired Alus’: either no Alu elements or no Alu elements flanking the exon within the window; ‘Non-complementary’: flanking Alus in the same orientation; ‘Complementary’: at least one sense Alu on one flank and at least one antisense Alu on the opposite flank. (**i**) Aggregate analysis of ILF2 and ILF3 binding ±400 bp around splice sites, comparing alternative exons spliced in control cells (gray) and ILF3-dependent alternative exons (pink). (**j**) RNA-seq tracks showing alternative splicing in *PILRB* and *ANAPC16*, with ILF3 eCLIP binding and Alu element locations. (**k**) Alternative splicing events in adult progenitor cells following *ILF3* knockdown (|ΔPSI| > 0.1, FDR < 0.05;). (**l**) Alternative splicing analysis in mouse ESCs comparing *Ilf2* knockout (data from^102^) and *Ilf2*/*3* knockdown (this study) (|ΔPSI| > 0.1, FDR < 0.05). (**m**) Differential analysis of splicing junctions between exons and transposable elements (JETs) after ILF3 degradation (blue: downregulated, red: upregulated; Padj < 0.01). (**n**) Repetitive element enrichment in differentially expressed JET transcripts (FC > 0, Padj < 0.01) relative to a randomized peak distribution. (**o**) Editing frequencies in differentially expressed JET transcripts after ILF3 degradation (n = 2 biological replicates; P values determined by Wilcoxon rank-sum test). (**p**) RT-qPCR validation of *ADAR* knockdown (n = 3 biological replicates; mean ± s.d.; P values determined by unpaired Student’s t-test). (**q**) Experimental design for JETs analysis after ILF3 degradation and *ADAR* knockdown. (**r**) JET events 24 hours after ILF3 degradation in control (left) and *ADAR* knockdown (right) cells (FC > 0, Padj < 0.01 in CTRL knockdown after 24 hours of ILF3 degradation). (**s**) Transcriptome changes after 24 and 96 hours of ILF3 degradation, highlighting transcripts with splice junctions between exons and transposable elements (JETs, Padj < 0.1). (**t**) RT-qPCR validation of doxycycline-inducible *UPF1* knockdown (n = 3 biological replicates; mean ± s.d.; P values determined by unpaired Student’s t-test).

**Extended Data Figure 8.**
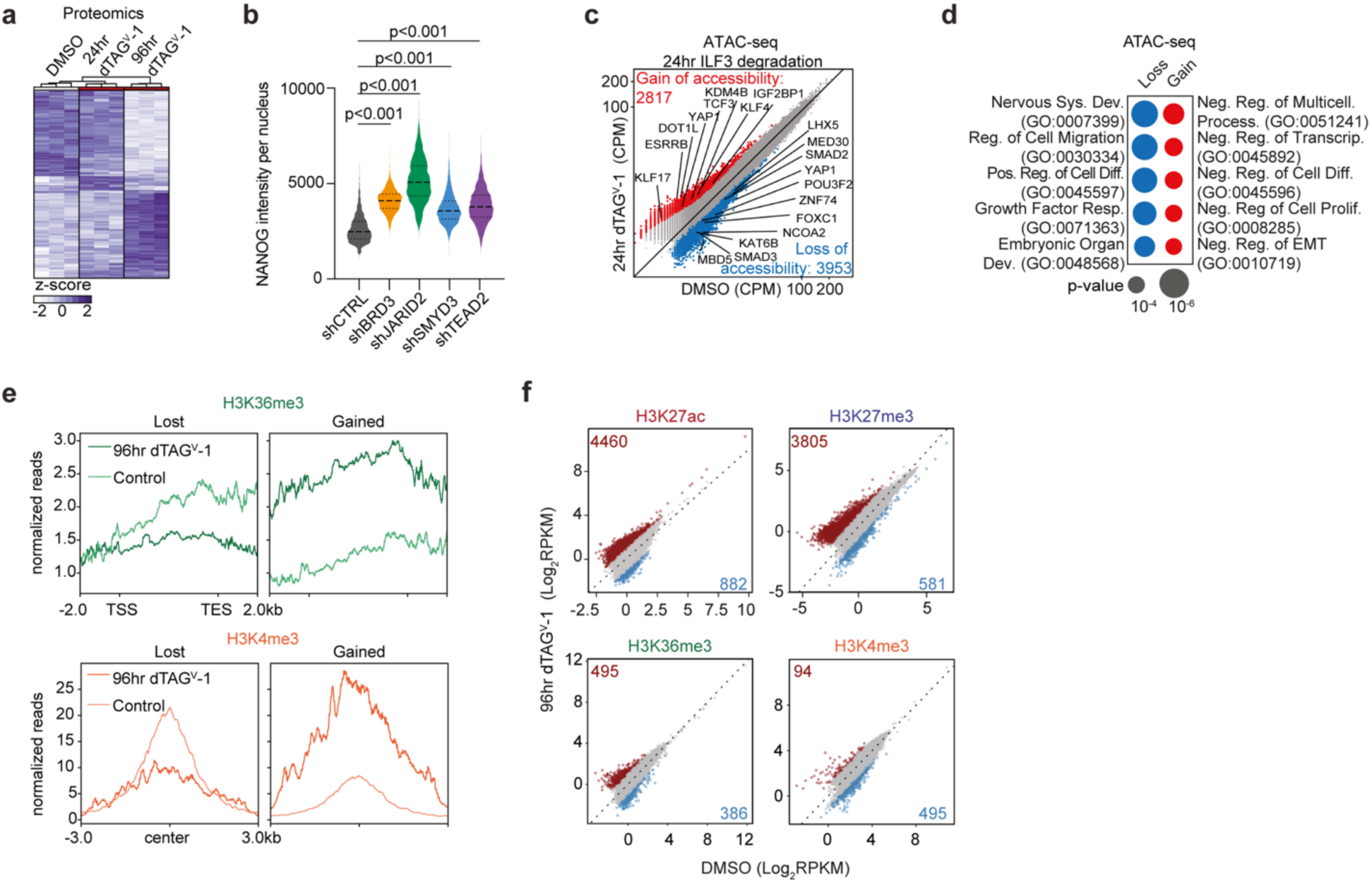
ILF3 depletion triggers extensive changes in the proteome and chromatin architecture. (**a**) Proteome changes after 0, 24, and 96 hours of ILF3 degradation in hPSCs (fold change > 1.2, P < 0.05). (**b**) Quantification of NANOG immunofluorescence intensity after knockdown of chromatin regulators in hPSCs under exit conditions by MAPK pathway inhibition (P values determined by one-way ANOVA with Dunnett’s multiple comparisons test). (**c**) ATAC-seq analysis after 24 hours of ILF3 degradation (n = 2 biological replicates; red: increased accessibility, blue: decreased accessibility; |FC| > 1.5, P < 0.05). (**d**) GO analysis of genes associated with regions showing altered chromatin accessibility (P values determined by two-tailed Fisher’s exact test). (**e**) Aggregate analysis of histone mark profiles after 96 hours of ILF3 degradation versus control (n = 3 biological replicates). (**f**) Differential analysis of histone modification patterns after 96 hours of ILF3 degradation (n = 3 biological replicates; RPKM>0.5; top 1% of peaks by area under the curve (AUC)).

